# Functional Characterization of Paillotin: An Immune Peptide Regulated by the Imd Pathway with Pathogen-Specific Roles in *Drosophila* Immunity

**DOI:** 10.1101/2025.05.12.653313

**Authors:** Yao Tian, Xiaojing Yue, Renjie Jiao, Mark A. Hanson, Bruno Lemaitre

## Abstract

Insects, such as *Drosophila* melanogaster, rely on innate immune defenses to combat microbial threats. Antimicrobial peptides (AMPs) play an important role in limiting pathogen entry and colonization. Despite intensive research into the regulation and biochemical properties of AMPs, their exact significance in vivo has remained uncertain due to the challenges of mutating small genes. Fortunately, recent technologies have enabled the mutation of individual AMP genes, overcome previous obstacles, and opened new avenues for research. In this study, we characterized one novel host-defense peptide, Paillotin (IM18, *CG33706*), using loss-of-function mutants. Paillotin is an ancient host defense peptide of Diptera, regulated by the Imd pathway. Loss of Paillotin does not impact the activity of either the Imd or Toll pathways. Importantly, we found that *Paillotin* mutants are viable but exhibit increased susceptibility to specific infections, particularly *Providencia burhodogranariea*. Paillotin was further found to contribute synergistically to defense against *P. burhodogranariea* when combined with other AMPs. However, we did not detect direct microbicidal activity of Paillotin *in vitro* in our hands. Taken together, our findings identify Paillotin as a novel host defense peptide acting downstream of Imd signaling, advancing our understanding of the *Drosophila* antimicrobial response.

## Introduction

Insects inhabit environments rich in microbes, necessitating a robust immune system to defend against opportunistic pathogens [1]. Unlike vertebrates, insects rely solely on innate immune defense mechanisms. The conservation of innate immunity across *Metazoans* and the powerful genetics of *Drosophila melanogaster* make this insect a powerful model for studying immune responses [2–4].

Flies exhibit behavioral immunity to prevent pathogens from entering their body cavity, such as grooming and avoidance of pathogens [5,6]. Once pathogens breach this barrier, immune responses are activated in different body regions to limit growth. Local immunity occurs at initial pathogen entry sites, like the gut or trachea. Systemic immunity involves cellular and humoral responses that take place in the body cavity upon systemic infection. Cellular responses involve hemocytes (blood cells), while humoral responses involve the production of host defense peptides as well as other effectors secreted by hemocytes and the fat body, an organ analogous to the mammalian liver [1,2]. Two NF-κB signaling pathways, the Toll and Imd pathways, regulate the humoral response. The principal effectors downstream of the Toll and Imd signaling pathways consist of AMPs, which are synthesized by both the fat body and hemocytes and then secreted into the hemolymph. AMPs are positively charged and can bind to the negatively charged membranes of microorganisms, disrupting membrane integrity through pore formation [1,7]. Upon infection, their expression is markedly increased to high levels in *Drosophila*, notably *Drosomycin* and *Diptericin,* which are often used as readouts for Toll and Imd activity, respectively [8]. Besides microbe killing, AMPs have been implicated in multiple processes such as tumor suppression [9,10], microbiota control [11], and neurodegeneration [12,13]. Currently, *D. melanogaster* is known to possess eight families of inducible AMPs, including antifungal peptides like Drosomycin, Baramicin, and Metchnikowin; Cecropins (four isoforms) exhibiting both antibacterial and antifungal activities *in vitro*; and Drosocin, Attacins (four isoforms), Defensin, and Diptericins (two isoforms), primarily displaying antibacterial activity [14–21]. Additionally, the *Drosophila* genome encodes numerous other host defense peptides, such as the Toll-regulated *Daisho* genes (two genes),

*Bomanins* (twelve genes), peptides encoded by *Baramicin A*, and the Imd-regulated peptide Buletin (product of the *Drosocin* gene), whose antimicrobial activity *in vitro* remains to be fully demonstrated, but functional studies have indicated their importance *in vivo* for surviving microbial infections [22–26].

Functional studies using single or combined mutations have demonstrated that these host defense peptides largely account for the critical roles of the Imd and Toll pathways in host survival following infection with Gram-negative bacteria, Gram-positive bacteria, or fungi. The antibacterial peptides Diptericin, Cecropin, Attacin, Drosocin, and Buletin account for most of the Imd pathway–mediated resistance to Gram-negative bacterial infection [25,27–29]. Drosomycin, Metchnikowin, Daisho (two genes), Baramicin A mediate much of the Toll-mediated resistance to fungal infection [23,24,26,27,30] , and Bomanins (twelve genes) appear to be the most important family of Toll-regulated host defense peptides contributing to resistance against Gram-positive bacteria and fungi [22,27,31]. These host defense peptides display both additive and synergistic effects, along with a high degree of specificity, whereby individual peptides greatly contribute to survival against a defined pathogen [27,29,32]. Some immune-induced peptides regulated by Toll and Imd have regulatory functions. This includes GNBP-like3 and Ibin A/B which have been reported to modulate Toll pathway activity, as well as Edin, which has been implicated in regulating hemocyte sessility [33–35].

Despite significant advances in understanding *Drosophila* host defense peptides, many peptides remain underexplored. The *Drosophila* genome encodes several other immune-induced peptides that remain poorly characterized, including notably Listericin, CG4269, and IM18 [1,36,37].

In this article, we have characterized the function of IM18, which we renamed Paillotin. We showed that Paillotin is an ancient host defense peptide regulated by the Imd pathway, which does not affect the regulation of the Imd or Toll pathways. Importantly, we found that *Paillotin* mutants exhibit increased susceptibility to specific infections, particularly *Providencia burhodogranariea*. Paillotin was also found to contribute synergistically to defense against *P. burhodogranariea* when combined with other AMPs, specifically Drosocin, Diptericins, and Attacins. However, we did not detect direct microbicidal activity of Paillotin *in vitro*. Taken together, our findings identify Paillotin as a novel host defense peptide downstream of Imd signaling, advancing our understanding of the *Drosophila* antimicrobial response.

## Results

### Paillotin is a conserved host defense peptide in *Diptera*

A previous proteomic analysis identified an immune inducible peptide called Immune-induced Molecule 18 (IM18) based on its mass by MALDI-TOF [38]. We rename the *IM18* gene (*CG33706*) as *Paillotin* (*Pai*) from André Paillot, who in 1920 discovered the humoral immune response of insects [39,40]. The gene encoding Paillotin was identified as *CG33706* in a region annotated as polycistronic with another gene, *CG10332*, making it challenging to distinguish their expression patterns (Figure 1*a*); the *CG10332-RA* transcript is identical to the *Pai-RA* transcript and contains the full coding sequence of both *CG10332* and *Pai*. Indeed, in FlyAtlas data, *IM18* and *CG10332* are shown with identical expression levels [41]. However, the characterization of IM18 as an immune-induced peptide pointed to the existence of two distinct gene products, perhaps regulated by the distinct transcripts *Pai-RB* and *CG10332-RA.* To confirm this, we monitored the expression of either *CG10332-RA* alone or the cumulative products of *CG10332-RA* and *Pai-RB* upon septic injury with either the Gram-negative bacterium *Escherichia coli* (*E. coli*) or the Gram-positive bacterium *Micrococcus luteus* (*M. luteus*). The results showed that the mRNA encoding *Pai-RB* (Figure 2*a*, *b*) but not *CG10332-RA* was strongly induced upon septic injury (Figure 2*c*, *d*), confirming that only *Pai-RB* is immune-induced at the transcriptional level.

**Figure 1:**
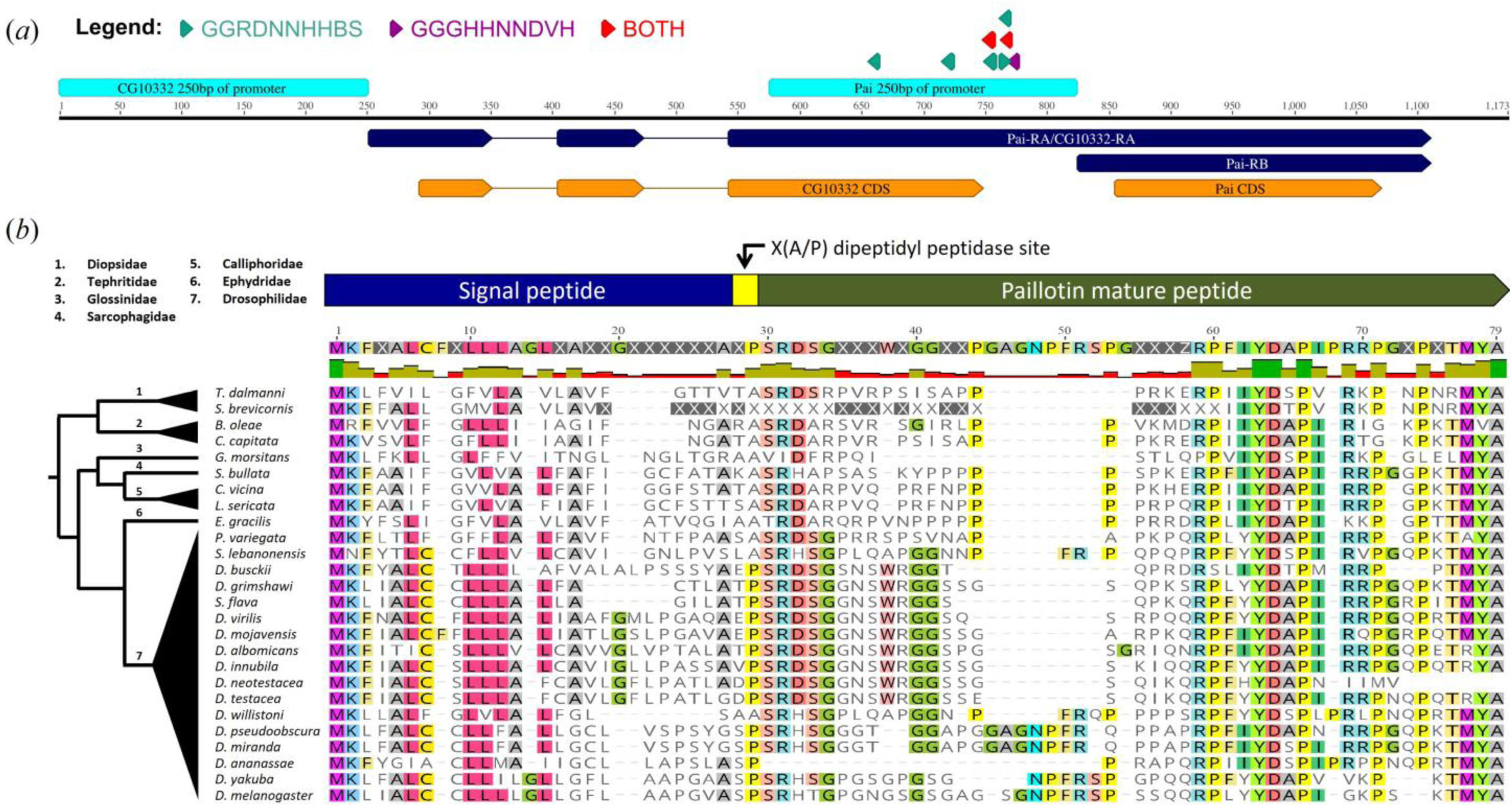
The *Paillotin* gene structure and conservation. (*a*) Schematic representation of the *Paillotin* (*Pai*) locus. The *Pai-RB* transcript is annotated as polycistronic, sharing a promoter and transcript with *CG10332* (*CG10332-RA*), indicating both gene CDS regions are co-transcribed. Promoter regions (250 bp upstream) for *Pai-RB* and *CG10332-RA* are shown in cyan. Colored triangles indicate putative NF-κB binding motifs: *Relish*-responsive motifs (GGRDNNHHBS) in teal, *Dif/Dorsal*-responsive motifs (GGGHNNNDVH) in purple. Red triangles represent sites containing overlapping matches to both motifs. (*b*) Sequence alignment of Paillotin orthologues across diverse *Brachyceran* flies revealed a conserved protein structure, including a signal peptide, a dipeptidyl peptidase cleavage site (XA/XP), and the mature Paillotin peptide. Highlighting corresponds to residues with >50% identity across the alignment.

**Figure 2:**
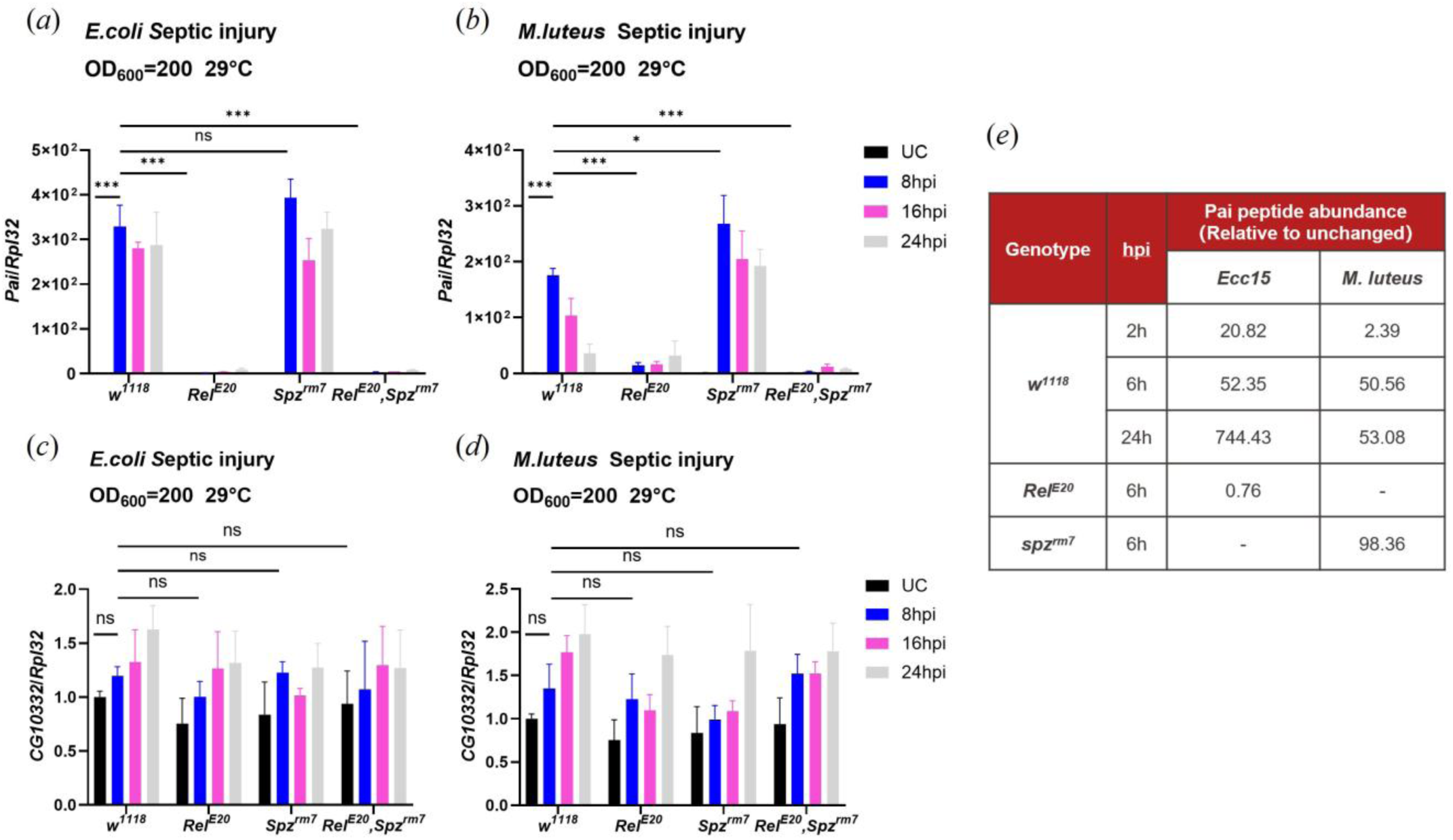
*Paillotin* is regulated by the Imd pathway. Transcript levels of *Paillotin*/*CG10332* in *w1118*, *RelishE20*, *spzrm7*, and *RelishE20*, *spzrm7* flies collected at 8 h, 16h, and 24h post-*E. coli* infection (*a*, *c*) and *M. luteus* infection (*b*, *d*). (*e*) Quantified Paillotin peptide levels of *w1118*, *RelishE20*, and *spzrm7* hemolymph upon either *Ecc15* or *M. luteus* infection (data extracted from (Rommelaere *et al.*, 2025)). Gene Expression was measured by qRT-PCR and normalized with *w1118* UC (unchallenged) flies set as a value of 1. The data shown in the figure are based on three independent experiments, with at least 30 flies per genotype at each time point in each experiment. Error bars represent SEM. Statistical significance was determined using two-way ANOVA (\**p*<0.05, ** *p*<0.01, *** *p*<0.001, n.s. = not significant, *p*>0.05).

The *Paillotin* gene encodes a 71-residue precursor peptide composed of a signal peptide, dipeptidylpeptidase motif, and mature peptide (Figure 1*b*). To investigate the evolutionary conservation of Paillotin, we used a recursive tBLASTn approach to screen outgroup fly species [42]. Orthologues of Paillotin were identified in *Diptera* families such as *Tephritidae* and *Diopsidae*, which share a common ancestor with *D. melanogaster* approximately 126 million years ago [43] (Figure 1*b*). The C-terminal domain displayed the highest level of conservation across species, suggesting a critical role in Paillotin function. The presence of conserved features, such as the two-residue dipeptidyl peptidase cleavage site (XA/XP) and a positive charge (+3.89 in *D. melanogaster*) commonly found in AMPs [24,44], supports the hypothesis that Paillotin plays a key role in host defense and could have microbicidal activities. This evolutionary conservation underscores Paillotin as an ancient host defense peptide in *Diptera*.

### *Paillotin* is regulated by the Imd pathway

To explore the expression pattern and regulation of *Paillotin* during systemic immune responses, we analyzed transcript levels in wild-type, Toll, and Imd pathway-deficient flies (Figure 2). Specifically, we challenged *w^1118^* (wild-type), *spz^rm7^* (deficient for the Toll pathway), *Relish^E20^* (deficient for Imd pathway) and *Relish^E20^*, *spz^rm7^* double mutant flies with either *E*. *coli,* an inducer of the Imd pathway (Figure 2*a*), or *M. luteus,* an inducer of the Toll pathway (Figure 2*b*). Expression of *Paillotin* was monitored at 8h, 16h, and 24h post-infection using qRT-PCR. We observed that *Paillotin* gene expression was strongly induced after both *M. luteus* and *E. coli* infections in *w^1118^* flies at all time points. However, this induction was completely abolished in *Relish^E20^* and *Relish^E20^*, *spz^rm7^* double mutants, but not in *spz^rm7^*flies, indicating *Paillotin* is regulated by the Imd pathway (Figure 2*a*, *b*). Consistent with this expression profile, we found multiple putative NF-κB responsive motifs within the first 200bp of promoter sequence upstream of gene transcription start sites of *Paillotin* [24,45] including five motifs uniquely matching the *Relish* consensus motif sequence (GGRDNNHHBS), one motif matching a *Dif/Dorsal* (GGGHHNNDVH) binding site, and two κB motifs with putative matches to either *Relish* and/or *Dif/Dorsal* binding sites. In contrast, no NF-κB binding motifs were identified in the 250bp upstream region of *IM18-RA/CG10332-RA*, consistent with its non-immune inducibility.

A recent proteomic analysis of *Drosophila* hemolymph following bacterial infection [46] indicated that Paillotin peptide was strongly induced in the hemolymph of wild-type flies infected with *M. luteus* or the Gram-negative bacterium *Pectobacterium carotovorum* (*Ecc15*), and this induction was abolished in *Relish^E20^* flies (Figure 2*e*). Consistent with its regulation by the Imd pathway, *Paillotin* was more induced upon infection with *Ecc15* than *M. luteus*. We conclude that Paillotin is a novel host defense peptide regulated at the transcriptional level by the Imd pathway and secreted into the hemolymph.

### Identification and characterization of *Paillotin* mutants

The *Drosophila* genome nexus represents a global sampling of the genetic diversity of *D. melanogaster* wild types subjected to genome sequencing [47]. Using a combination of the genome nexus and more detailed variant information in the *Drosophila* Genetic Reference Panel (DGRP) [48], we identified two strains from Kenya (KR7, KR4N) encoding a premature stop codon in *Paillotin* (“Gly37⁕”, numbering per start codon). Globally, we could assess the *Paillotin* region in 789 genome-sequenced strains, finding 39 that encode putative loss-of-function mutations (Table S1). The majority of these mutant strains were collected from Winters California and Raleigh North Carolina, which have a high frequency of putative loss-of-function mutations (∼26% and ∼8% respectively within these populations). We characterized two of the loss-of-function mutations segregating in the DGRP further (Figure 3*a*). The first mutation (2R_19488505_DEL) is found in 17 genotypes from the DGRP, including DGRP-41, which deletes 61 nucleotides within the Paillotin mature peptide coding region, leading to a frameshift and protein truncation (referred to as *Pai*^△^*^41^*). The second mutation (2R_19488690_SNP), found in DGRP-370 (*Pai* ^△^*^370^*), involves a C-T transition that disrupts the *Paillotin* start codon. We utilized these naturally-occurring mutations to study the role of *Paillotin* in host defense by backcrossing *Pai*^△^*^41^* and *Pai*^△^*^370^* into the *w^1118^* iso DrosDel background for seven generations.

**Figure 3:**
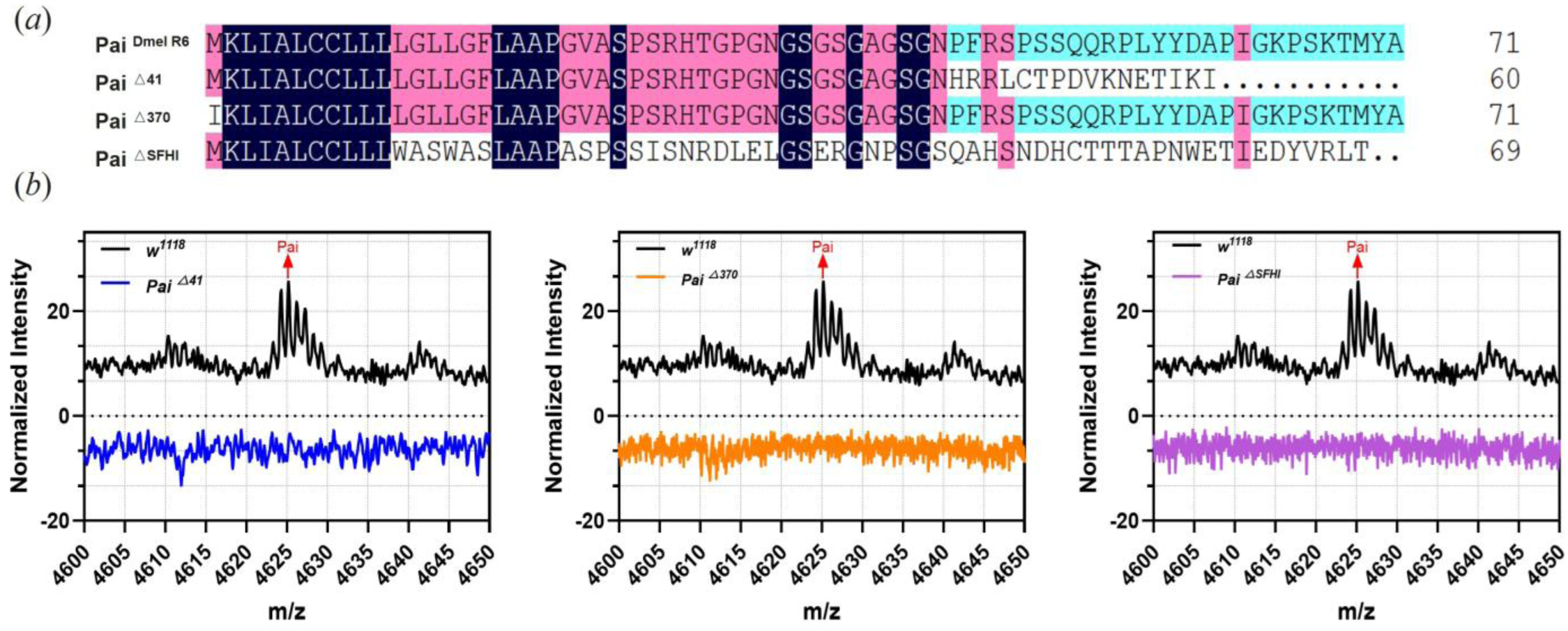
Description and validation of *Pai*△*41*, *Pai*△*370*, and *Pai*△*SFHI* mutations. (*a*) Multiple sequence alignment between the Dmel_R6 reference genome Paillotin and Paillotin peptide variants. Conserved residues are highlighted by color coding according to sequence similarity. (*b*) MALDI-TOF proteomic analysis of hemolymph from *w1118*, *Pai*△*41*, *Pai*△*370*, and *Pai*△*SFHI* male flies collected 24 hours after septic injury with *Ecc15*.

In addition, we used a third mutant referred to as *Pai*^△^*^SFH^*, a gift from the Guangzhou Drosophila Resource Center (Figure 3*a*), which was generated by CRISPR-Cas9 and involves a 20-nucleotide deletion that causes a complex frameshift, resulting in a nonsense peptide. The precise nucleotide alterations associated with each *Paillotin* mutant allele used in this study are presented in Supplementary Figure S1. For *Pai*^△^*^SFHI^*, only the *X* chromosome was replaced by the *w^1118^* iso DrosDel *X* chromosome.

RT-qPCR confirmed reduced *Paillotin* gene expression in *Pai*^△^*^41^* and *Pai*^△^*^370^*, while *Paillotin* expression was completely absent in *Pai*^△^*^SFHI^* (Figure S2). We next performed a MALDI-TOF analysis of hemolymph from adult flies infected with *P. c. carotovara Ecc15*. The results revealed the absence of the 4625Da Paillotin peaks in *Pai*^△^*^41^*, *Pai*^△^*^370^*, and *Pai*^△^*^SFHI^* flies (Figure 3*b*). All the mutants were viable without morphological defects.

### *Paillotin* does not regulate the Imd and Toll pathways

Some host defense peptides, such as mammalian LL37 or β-Defensins, have been shown to function similarly to cytokines, modulating immune system activity [49,50]. This prompted us to assess whether Paillotin could influence Toll and Imd pathway activity. To address this, we monitored the expression of *Diptericin A* (a readout of the Imd pathway) in wild-type (*w^1118^*) and *Paillotin* -deficient flies (*Pai*^△^*^41^* and *Pai*^△^*^370^*) at 6- and 12-hours post-infection with *Ecc15* and *Drosomycin* (a readout of the Toll pathway) at 8- and 24-hours post-infection with *M. luteus* (Figure 4*a*, *b*). The expression levels of *Diptericin A* in *Pai*^△^*^41^* and *Pai*^△^*^370^* mutants were comparable to those in wild-type flies at both time points, indicating no effect of *Paillotin* on Imd pathway activity. Similarly, the expression of *Drosomycin* in *Pai*^△^*^41^* and *Pai*^△^*^370^* mutants showed no significant difference compared to *w^1118^*, suggesting that *Paillotin* is not involved in regulating the Toll pathway. Together, these results demonstrate that *Paillotin* is not a regulator of the Imd or Toll pathways.

**Figure 4:**
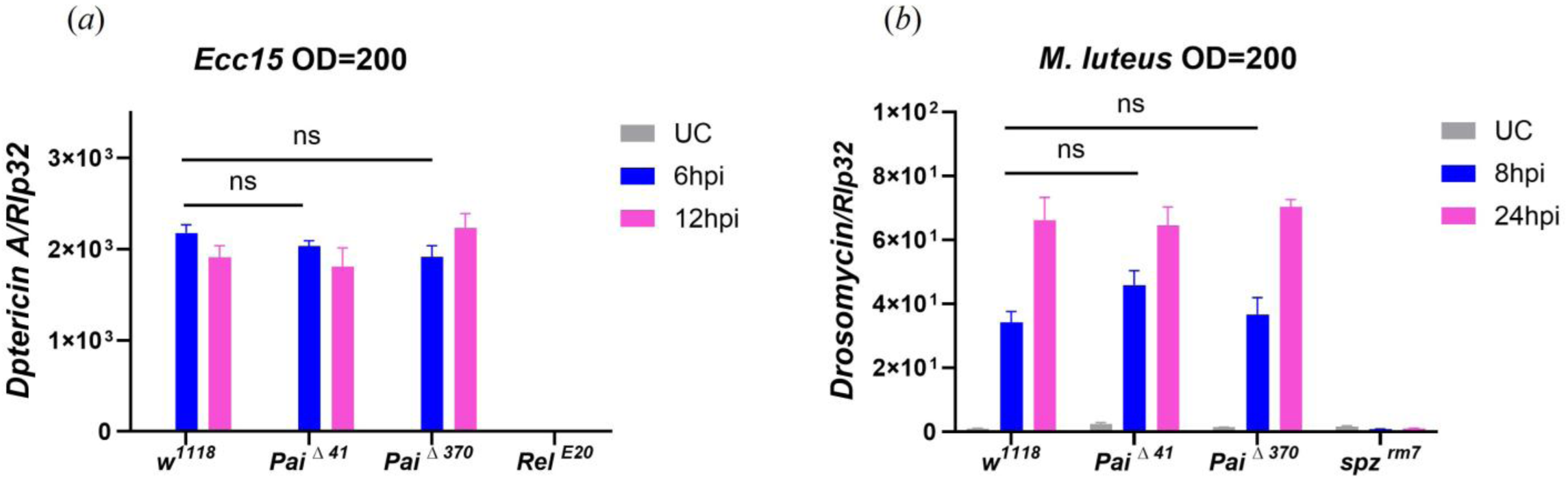
Paillotin does not impact the Imd or Toll pathways. The expression of *Diptericin A* (*a*) at 6 or 12h post-infection with *Ecc15* and *Drosomycin* (*b*) at 8h or 24h post-infection with *M. luteus* were monitored in *w1118*, *Pai*△*41*, and *Pai*△*370* flies. *RelishE20* and *spzrm7* were used as the positive control. Expression was normalized to *w1118* UC (unchallenged) flies set as a value of 1. The data shown in the figure are based on three independent experiments, with at least 30 flies per genotype at each time point in each experiment. Error bars represent SEM. Statistical analysis was performed using two-way ANOVA. For *Diptericin A*, significant effects were detected for genotype (F(3,24) = 148.1, p < 0.0001), time (F(2,24) = 332.8, p < 0.0001), and their interaction (F(6,24) = 38.98, p < 0.0001). For *Drosomycin*, significant effects were also observed for genotype (F(3,24) = 72.82, p < 0.0001), time (F(2,24) = 191.5, p < 0.0001), and interaction (F(6,24) = 22.84, p < 0.0001). However, Tukey’s post hoc comparisons revealed no significant difference between *w1118* and *Paillotin* mutants at any time point (n.s. = not significant).

### *Paillotin* mutant flies are susceptible to *P. burhodogranariea* infection

Recent studies have demonstrated that the deletion of a single AMP gene can significantly increase susceptibility to specific pathogens. For instance, the *Diptericin A* mutation leads to marked susceptibility to *Providencia rettgeri*, while the *Diptericin B* mutation greatly reduces resistance to *Acetobacter sp.* [29,32]. To investigate the role of Paillotin in resistance to infections, we examined its contribution against a range of Gram-negative, Gram-positive bacteria, and fungi. In these experiments, we compared the survival of *Pai*^△^*^41^*, *Pai*^△^*^370^*, and *Pai*^△^*^SFHI^* flies to that of the wild-type (*w^1118^*), *Relish^E20^, Bom^△55^,* and *spz^rm7^* immune-deficient mutants. Flies were systemically infected by bacteria using a needle dipped in a bacterial pellet, a mode of infection that causes cuticle injury. To exclude a role of *Paillotin* in resistance to injury, we first monitored the susceptibility of *Paillotin* mutants to clean injury, and did not detect any susceptibility after clean injury alone (Figure S3). Regarding Gram-negative bacteria, there was no statistically significant difference in survival between wild-type and *Paillotin* mutants following infections with *P. rettgeri*, *Ecc15*, or *Enterobacter cloacae* (Figure S4*a*-*c*). However, all three *Paillotin* mutant flies (*Pai*^△^*^41^*, *Pai*^△^*^370^*, *Pai*^△^*^SFHI^*) exhibited a modest but significant increase in susceptibility to septic injury with *Providencia burhodogranariea* (*P*. *burhodogranariea*) (Figure 5*a*). This susceptibility was less marked than *Relish^E20^* flies, indicating that other factors downstream of the Imd pathway contribute to survival upon *P. burhodogranariea* infection. In addition, we observed higher *P. burhodogranariea* bacterial load in *Paillotin* mutants compared to wild-type, indicating that Paillotin suppresses bacterial growth (Figure 5*b*). For Gram-positive bacteria, *Paillotin* mutants showed no consistent increased susceptibility to *Enterococcus faecalis* (Figure 5*c*), *Bacillus subtilis*, *Listeria monocytogenes*, or *Staphylococcus aureus* infections (Figure S4*d*-*f*). Similarly, fungal infections with *Beauveria bassiana* (Figure 5*d*), *Metarhizium anisopliae*, or *Metarhizium robertsii* did not reveal any increased susceptibility of *Paillotin* mutants compared to wild-type (Figure S4*g*-*h*).

**Figure 5:**
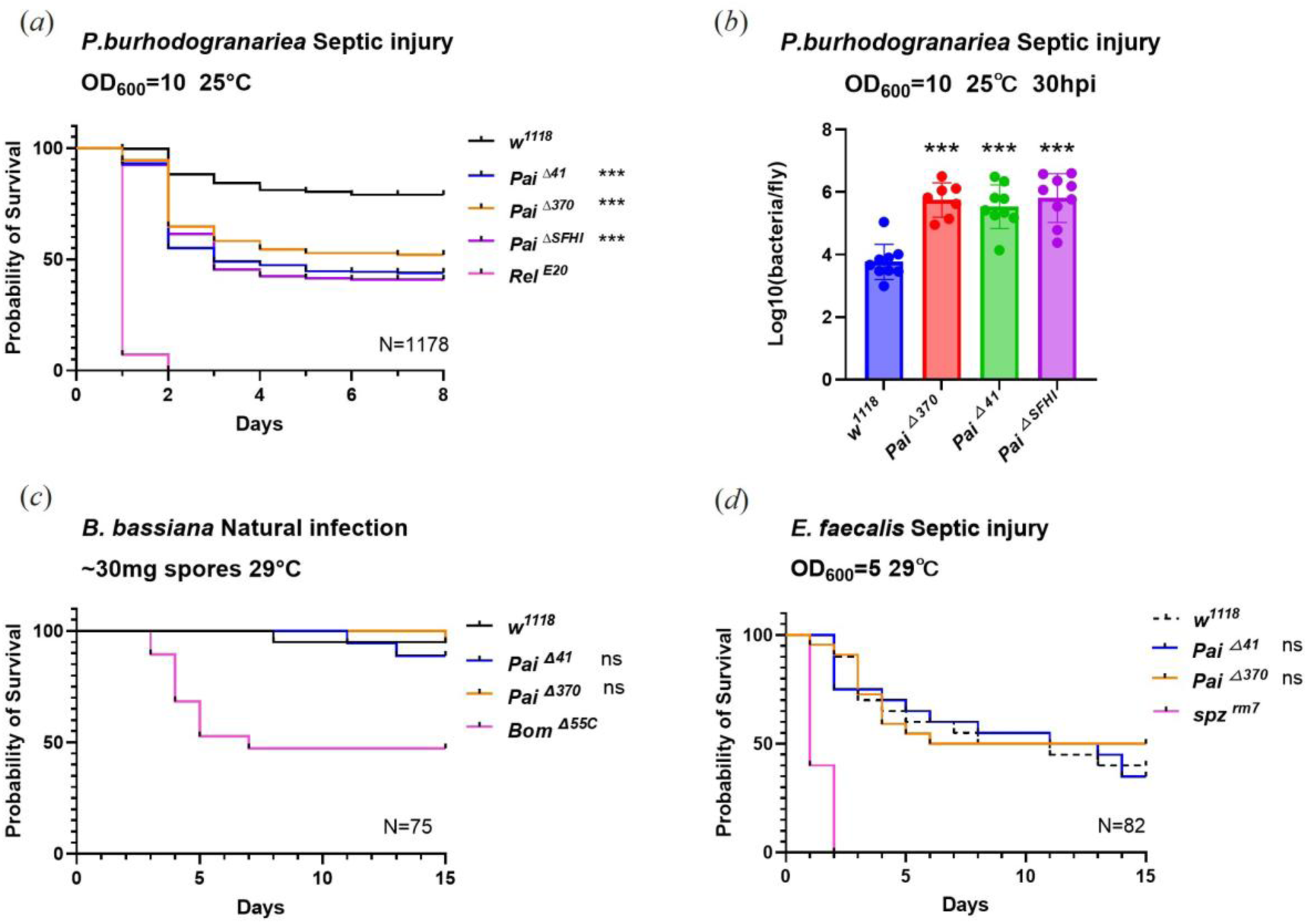
*Paillotin* mutants are susceptible to *P. burhodogranariea* infection. (*a*) The three *Paillotin* mutants exhibit increased susceptibility following septic injury with *P. burhodogranariea* at the indicated bacterial inoculum. (*b*) Increased bacterial load in *Paillotin* mutants at 30 hpi. Bacterial load was log₁₀-transformed prior to statistical analysis. One-way ANOVA revealed a significant genotype effect (F(3,30) = 18.95, p < 0.0001). Post hoc Dunnett’s test confirmed that all three mutant lines carried significantly higher bacterial loads than *w1118* (p < 0.0001 for all comparisons). Data points represent the mean bacterial load per fly, derived from individual experiments using pools of four flies per genotype following *P. burhodogranariea* infection. No consistent effect of the *Paillotin* mutation on resistance to the Gram-positive bacterium *Enterococcus faecalis* (*c*) or the fungus *Beauveria bassiana* (*d*). Survival data were analyzed using the Kaplan–Meier method. Statistical significance is indicated relative to the *w1118* (*** *p*<0.001, n.s. = not significant, *p*>0.05). N=total number of flies in experiments.

### No *in vitro* activity of Paillotin peptide

Cecropins were the first inducible AMPs identified in moths [14]. *In vitro* studies have shown that Cecropins are highly effective against a wide range of Gram-negative bacteria at concentrations lower than those induced in insects during infection (25–50 µM) [51]. To investigate the antimicrobial potential of Paillotin *in vitro*, we produced the peptide and evaluated its activity against *P. burhodogranariea* at 100µM alone or in combination with sublethal concentrations of Cecropin A. However, Paillotin exhibited no detectable microbicidal activity against *P. burhodogranariea in vitro* (Figure S5*a*). It also failed to inhibit the growth of other Gram-negative bacteria, including *Ecc15* and *E. coli* (Figure S5*b*, *c*). Thus, we found no activity of Paillotin *in vitro* under these conditions.

### Paillotin synergistically contributes to the defense against *P. burhodogranariea* with other AMPs

Recent studies have demonstrated synergistic or additive cooperation among AMPs, notably against *P. burhodogranaria*. While single mutations in *Drosocin*, *Attacins*, or *Diptericins* have minimal overall effect, a compound mutant referred to as the “*Group B”* AMP mutant lacking *Drosocin*, *Attacins A, B, C* and *D*, and *Diptericins A* and *B* [full genotype: *w^1118^; AttC^Mi^, Dro-AttAB^SK2^, Dpt^SK1^; AttD^SK1^*], displays marked susceptibility to *P. burhodogranariea* infection [27]. To investigate potential interactions between *Paillotin* and these AMPs, we used genetic crosses to combine the *Paillotin* mutation with *Group B* mutants. We initially infected *Group B* mutants, *Paillotin* mutants, and *Group B, Paillotin* compound mutants [full genotype: *w^1118^; AttC^Mi^, Dro-AttAB^SK2^, Dpt^SK1^, Pai*] with *P. burhodogranariea*. However, we observed no synergistic or additive effects, as the introduction of *Paillotin* mutations into *Group B* mutants did not further increase susceptibility. This lack of an observable effect may be due to the already high susceptibility of *Group B* AMP mutants to this bacterium, even at a low inoculum (OD600=0.1), potentially masking any additional contribution from the *Paillotin* mutation (Figure 6, dashed curves). To circumvent this, we assessed the impact of *Paillotin* homozygous deficiency in flies heterozygous for *Group B* AMP mutations (*Group B/+*), which depresses the inducibility of AMPs compared to homozygotes [27]. Notably, in this sensitized genetic background, the absence of *Paillotin*, regardless of the allele used, significantly increased susceptibility of *Group B/+* flies to *P. burhodogranariea* (Figure 6*a*-*c*, solid curves). This co-occurring loss of resistance reflects a synergistic effect on the Hazard Ratio (HR), evidenced by increased mortality beyond additive effects (*Paillotin* alone, HR =-0.17, *p* =0.01; *Group B* alone, HR =-1.06, *p*<0.001; *Paillotin*\**Group B* interaction, HR =-0.15, *p*=0.02). In addition, a marginally significant trend consistent with a synergistic effect between *Paillotin* and *Group B* mutants was also seen for infections with the Gram-negative bacteria *Providencia rustigianii* and *Providencia vermicola*, but not *E. coli* (Figure S6). This supports the notion that Paillotin confers a fairly specific resistance to *P. burhodogranariea*, perhaps also extending to other *Providencia* species.

**Figure 6:**
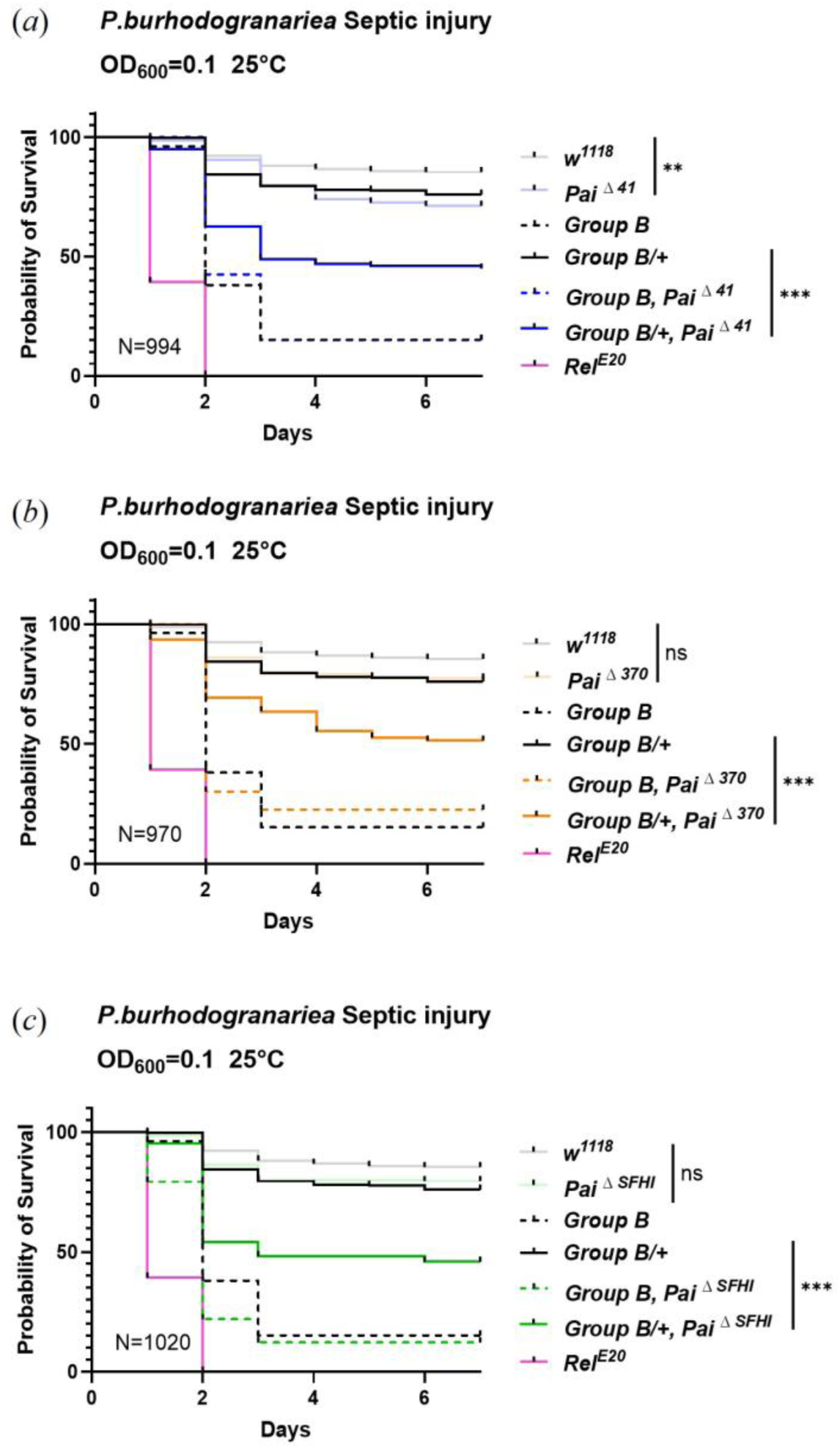
Paillotin synergistically contributes to defense against *P. burhodogranariea* with other Imd-regulated AMPs. (*a-c*) Survival curves showing the synergistic effects between different *Paillotin* mutants (*Pai*△*41*, *Pai*△*370*, *Pai*△*SFHI*) and *Group B* mutations (*Drosocin*, *Attacin A, B, C,* and *Diptericin A, B*) in the heterozygous background (*Group B/+*). Survival curves were analyzed using the Cox proportional hazards (CoxPH) model in R version 3.6.3. (** *p*<0.01, *** *p*<0.001, n.s. = not significant, *p*>0.05). N=total number of flies in experiments.

### Overexpression of Paillotin restores resistance in Paillotin-deficient flies against *P. burhodogranariea*

The observation that three distinct null mutations in *Paillotin* are associated with a higher susceptibility and the synergistic effect between Paillotin and other AMPs against infection provide strong evidence for a role of this host defense peptide in resisting *P. burhodogranariea*. To reinforce this finding, we performed a genetic rescue experiment by combining the *Paillotin* mutation with a *UAS-Pai* transgene. We initially observed that overexpression of *Paillotin* in a wild-type background was not sufficient to enhance resistance to *P. burhodogranariea* (Figure S7). In contrast, overexpression of *Paillotin* using either the *C564-Gal4* (a Gal4 driver with strong expression in the fat body) or *Actin-Gal4* (ubiquitous) driver restored pathogen resistance to near wild-type levels in the *Pai*^△41^ (Figure 7*a*) and *Pai*^△370^ (Figure 7*b*) mutant backgrounds. The observation that three distinct *Pai* alleles confer susceptibility, together with the results of the rescue experiment, collectively demonstrates that Paillotin acts as a pathogen-specific host defense peptide essential for restricting *P. burhodogranariea* proliferation.

**Figure 7:**
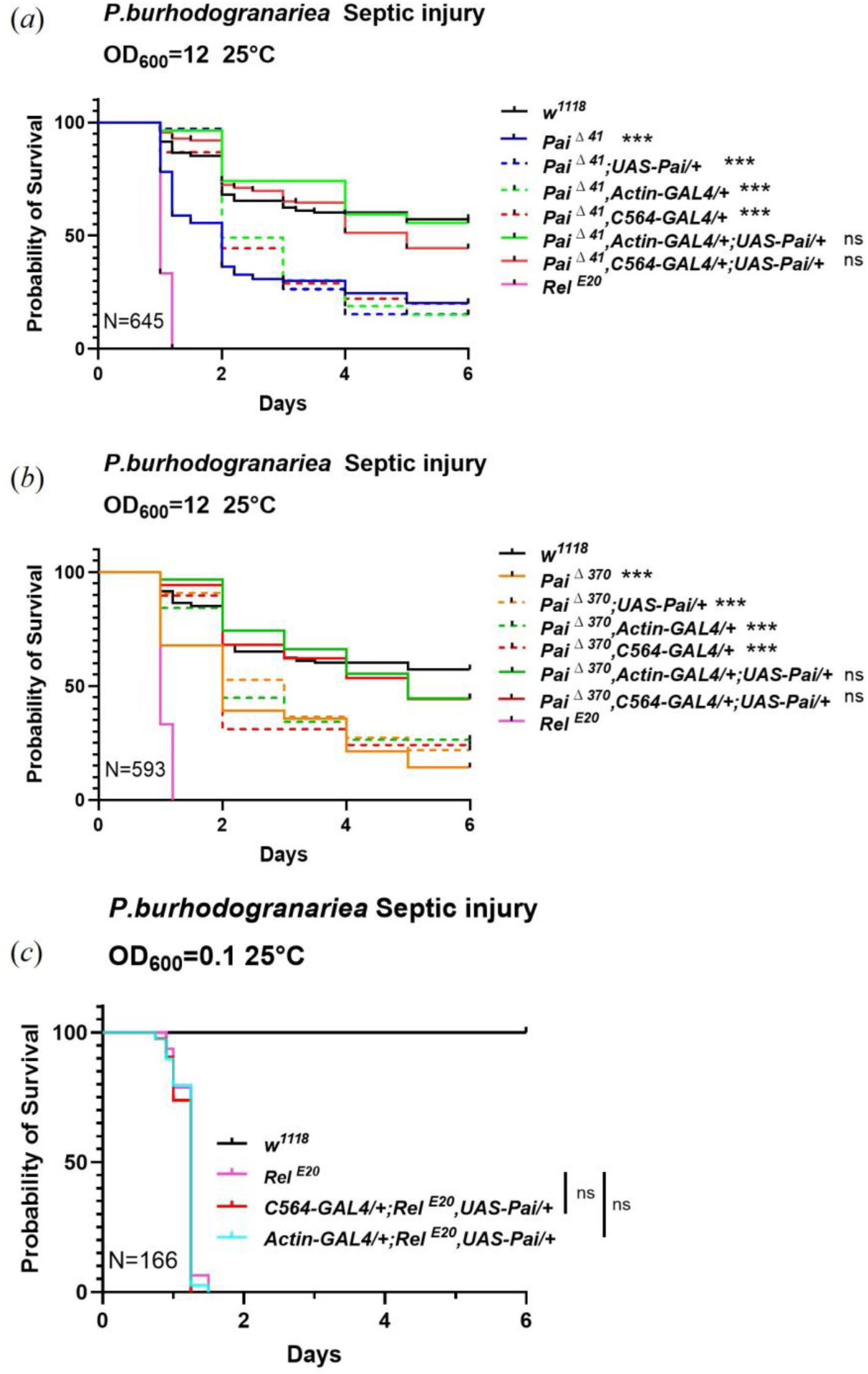
Over-expression of *Paillotin* rescues the resistance of *Paillotin*-deficient flies against *P. burhodogranariea*. (*a-b*) Overexpression of *Paillotin* via a combination of the *C564-Gal4* or *Actin-Gal4* and *UAS-Pai* constructs enhances the resistance of the Paillotin mutant to *P. burhodogranariea* infection. (*c*) Overexpression of *Paillotin* in the *RelishE20* background failed to restore resistance. Survival curves were analyzed using the Kaplan-Meier method. (*** *p*<0.001, n.s. = not significant, *p*>0.05). N=total number of flies in experiments.

The Imd and Toll signaling pathways are essential for insect immunity, with double pathway-deficient flies exhibiting extreme susceptibility to infection due to their inability to upregulate hundreds of immune effector genes, particularly AMPs [52]. Previous work has demonstrated that transgenic overexpression of individual AMPs in some cases is sufficient to restore resistance in *Imd, Toll-deficient* flies [53]. Building on this paradigm, we generated flies overexpressing *Paillotin* using *Actin-Gal4* or *C564-Gal4* drivers in *Relish^E20^* mutant flies (Figure 7*c*). However, *Paillotin* overexpression failed to confer protection against *P. burhodogranariea* infection. While we cannot exclude that Paillotin was not expressed at a sufficient level to overcome the extreme susceptibility of *Relish* mutants, this suggests that *Paillotin* alone is insufficient and the host requires the presence of other AMPs for the defense response to function effectively, consistent with the synergistic interactions observed with *Group B/+* heterozygous mutants.

## Discussion

In this study, we characterized Paillotin, an evolutionarily conserved host-defense peptide regulated by the Imd pathway. Use of three *Paillotin* mutants reveals a role of *Paillotin* in the resistance to *P. burhodogranariea* infection, and perhaps other moderately virulent *Providencia* species, without altering survival to other tested bacterial or fungal species. Furthermore, the *Paillotin* mutation enhances the susceptibility to *P. burhodogranariea* in flies heterozygous for mutations in *Drosocin*, *Diptericins A and B*, and *Attacins A, B, and C.* Taken together, the consistent susceptibility of three independent *Paillotin* alleles to *P. burhodogranariea*, accompanied by elevated bacterial loads, the resistance of *Paillotin*-deficient flies to other tested pathogens, and the successful rescue of the phenotype by overexpression, all provide strong evidence that *Paillotin* functions as a host defense peptide specifically effective against this bacterium. A high degree of specificity has been observed for other host defense peptides, whose inactivation frequently causes pathogen-specific immune deficiency. This has been illustrated not only for Diptericin A and B as described previously [29,32], but also for Drosocin, which provides significant protection against *E. cloacae* [25,27]. The existence of such specificity between a defined host defense peptide and particular pathogens suggests that these genes have been evolutionarily maintained under selective pressure imposed by specific microbes encountered in natural environments [54].

Previous studies have shown that resistance to *P. burhodogranariea* involves multiple antimicrobial peptides, including Buletin (encoded by the *Drosocin* gene), Diptericins A and B, and Attacins A, B, C, and D. While single AMP mutations in many genes do not greatly impact survival following infection with *P. burhodogranariea*, compound mutants lacking *Drosocin*, *Diptericins* and *Attacins* exhibit increased susceptibility, suggesting the presence of a synergistic effect among these peptides [27]. More recently, the absence of Buletin—a peptide generated by furin cleavage of the Drosocin precursor—results in notable survival differences following *P. burhodogranariea* infection [25]. Thus, host resistance to *P. burhodogranariea* is mediated by the collective action of multiple host defense peptides, and our study now adds Paillotin to this list. Future studies may further reveal additional roles of Paillotin, either alone or in combination with other host defense peptides, in resistance to other pathogens.

We were unable to detect any direct *in vitro* microbicidal activity of Paillotin against *P. burhodogranariea* and other bacteria, either alone or in combination with Cecropin. This reflects a common mismatch between in vitro and in vivo activities. Despite demonstrating effective antimicrobial activity in vivo in relevant animal models, many AMPs show minimal or no activity in standard minimal inhibitory concentration (MIC) or minimal microbicidal concentration (MMC) assays when tested under physiological salt concentrations or in serum-containing environments [55–58]. Several factors could explain this lack of detectable activity: i) Paillotin may depend on host-specific factors for its function, such as the requirement for a cofactor or modulation by environmental cues (e.g., pH or ionic strength), or ii) synergistic interactions with other immune molecules that are only present *in vivo*. Similar synergistic mechanisms have been observed in other systems. For instance, in *Galleria mellonella* (the greater wax moth), an anionic antimicrobial peptide (anionic peptide 2) enhances lysozyme-mediated membrane perforation, facilitating the elimination of *E. coli* [59]. Furthermore, AMPs often enhance each other’s efficacy through complementary mechanisms of action [21,60]. For instance, the bumblebee linear peptides hymenoptaecin and abaecin exhibit a synergistic interaction against *E. coli*. Although abaecin alone shows no detectable activity at concentrations up to 200 μM, hymenoptaecin disrupts the bacterial membrane, allowing abaecin to enhance bactericidal effects at doses as low as 1.25 μM [61]. In *Tenebrio molitor*, RNAi-mediated knock-down of the defensin Tenecin-1 did not affect host survival but reduced bacterial load following *S. aureus* infection. However, simultaneous knock-down of Tenecin-1 with other AMPs, such as the coleoptericin Tenecin-2 or the attacin Tenecin-4, resulted in increased mortality and higher bacterial loads, highlighting functional complementarity among AMPs during infection [62]. In vitro experiments showed that Attacin, an antibacterial protein that accumulates in the hemolymph of the giant silk moth *Hyalophora cecropia*, compromises the permeability barrier function of the *E. coli* outer membrane. Pretreatment with Attacin significantly increases the sensitivity of *E. coli* to Cecropin B, another antimicrobial protein present in the hemolymph of *H. cecropia* [63]. In addition, recent studies have pointed to the role of host defense peptides in resilience mechanisms, including tolerance to host immune molecules [46] and pathogen toxins [26,31]. However, we did not detect a role for Paillotin in resilience to the fungal toxins Destruxin A or Verruculogen (Figure S8). Interestingly, in contrast to recent studies [26,31], we did not detect a susceptibility of Toll-deficient *spz^rm7^* flies to these two toxins (Figure S8). Of note, the description of DIMs has been done largely by collecting peptides in Trifluoroacetic acid, which is known to modify peptides by, for example, deamidating NG motifs [24,64]. It is possible that the endogenous peptide products of the host immune response differ from the peptides observed in proteomic analysis and synthetic peptides used in recent studies, in unknown ways. Early studies on AMPs typically found striking activity *in vitro* using peptide extracts purified from endogenous hemolymph or cell line supernatant [14,17,19,21,65]. Future studies may benefit from a return to such approaches, including gel overlay experiments that can simplify the isolation of the activity of single peptides from endogenous samples [66] or collecting crude peptide extracts from overexpressing cells or larval hemolymph. However, future studies might also explore whether Paillotin protects against virulence factors specific to *P. burhodogranariea*. For now, despite displaying many features of AMPs, we cannot conclusively classify Paillotin as a bona fide AMP. It should be classified as a host defense peptide until its molecular mode of action is fully determined.

The use of compound module-deficient mutants has revealed the major role of the Imd pathway, and to a lesser extent, melanization, but not the cellular response to survive *P. burhodogranariea* infection [67]. Since Paillotin does not affect Toll and Imd pathway activities, we cannot exclude a role of this peptide in the melanization reaction. However, this role would have to be subtle as *Paillotin* mutants display a wild-type blackening reaction upon clean injury (Figure S9*a*). Previous studies have shown that flies lacking 14 AMPs exhibit increased microbiota abundance and altered composition in old flies, suggesting that AMPs are involved not only in pathogen resistance but also in shaping the environmental microbiota [11]. However, we did not detect any significant differences in gut microbiota load between wild-type flies and *Paillotin* mutants (Figure S9*b*).

Interestingly, there is a relatively high frequency of loss-of-function mutations segregating in wild *Drosophila* populations (Table S1), suggesting Paillotin is not integral to the generic host defense. This observation is consistent with our results showing that the *Paillotin* mutation affects defense against only a few bacteria out of the pathogens tested, suggesting it is dispensable in defense against many microbes, but useful against some microbes like *Providencia sp.* whose presence is stochastic in wild-caught flies [68].

To conclude, our study characterizes a new *Drosophila* host defense peptide, Paillotin, which plays a specific role against *P. burhodogranariea*. IM18/Paillotin is the last gene encoding an immune molecule identified by MALDI-TOF in the 1998 study [38] to be characterized, following the successful characterization of Bomanins [30], Daisho [23], Buletin [25], and Baramicin [24,26]. Together, these studies and the present article provide important insights into the *Drosophila* humoral immune response, revealing a more complex picture than initially anticipated. Using the power of *Drosophila* genetics to dissect how immune effectors individually or collectively contribute to host defense provides a general principle on the architecture of innate immune systems.

## Materials and Methods

### Sequence Comparisons

Genomic sequences were obtained from GenBank default reference assemblies [69], from the DPGP via the PopFly website (www.popfly.uab.cat)[47], and DGRP sequence data from the DGRP database [48] (http://dgrp2.gnets.ncsu.edu/). Sequence comparisons and alignment figures were prepared using Geneious R10 [70]. Putative NF-κB binding sites were annotated using the *Relish* motif “GGRDNNHHBS” as described in Copley et al [45], along with a manually curated amalgam motif “GGGHHNNDVH” described in Hanson et al [24].

Paillotin annotation across species was performed as described in Hanson et al [42]. In brief, a recursive tBLASTn approach was used with an artificially high E-value (E < 1000) to compensate for the short length of the AMP query sequence. All the best BLAST hits were assessed manually. If no Paillotin was found in a genome or lineage, only the conserved C-terminus was used as a query in a repeat attempt (improving sensitivity). No *Paillotin* candidates were discovered outside *Brachycera*, which included screens of the sequenced nematoceran genomes of various mosquitoes, *Chironomus riparius, Clogmia albipunctata, Tipula unca, Lutzomyia longipalpis, Phlebotomus papatasi, Sylvicola cinctus,* and *Bibio marci*.

### Fly Strains

Flies were maintained at 25°C on standard cornmeal-agar media. *w^1118^*flies were used as wild-type controls. The positive controls for infection assays for Gram-positive and Gram-negative infections were *spz^rm7^* and *Relish^E20^*. The *Pai*^△^*^41^* and *Pai*^△^*^370^* mutations are loss-of-function variants in the *IM18* gene within the DGRP, and *Pai*^△^*^SFHI^* was generated via CRISPR/Cas9. The guide RNAs that targeted *Pai* were cloned into the pUAST-attB plasmids before injecting into the *y [1] M {RFP [3xP3.PB] GFP[E.3xP3] =vas-int.Dm}ZH-2A w [*]; M{3xP3-RFP.attP} ZH-86Fb* (BL24749) to get gRNA transgenic flies. The gRNA flies were then crossed with flies containing nos-Cas9 to get mutants. The *UAS-Pai* constructs were prepared using the TOPO pENTR entry vector and cloned into the pTW system. Genetic variants were backcrossed into the DrosDel isogenic background over seven generations as previously described [71] using the following primers that are specific to an indel in the *Pai* locus common to both the *Pai*^△^*^41^* and *Pai*^△^*^370^*, but which is absent in the DrosDel background: Pai_41F (ACGTACGGTCAAAAGCAACTACTTCG) and Pai_R3mut1 (CCAATCCCAAGGGCACTCGTATATG), amplified with a 66°C annealing temperature. All the fly strains used in the paper are listed in Table S2.

### MALDI-TOF Proteomic Analysis

Hemolymph samples were collected from *Ecc15*-infected flies (OD600 = 200, 24 hours post-infection, 50 flies/sample) using a nanoinjector (Nanoject, Drummond Scientific, Broomall, PA) for MALDI-TOF analysis. Trifluoroacetic acid (0.1% TFA) was added to the hemolymph before mixing with the acetonitrile matrix. Samples are processed by SPE (Zip-Tip C18) before MALDI. Peaks were identified based on corresponding m/z values from previous studies [24,72] in ISIC’s Mass Spectrometry facility of EPFL. Spectra were visualized using Origin software, with figures prepared in Prism 10.

### Gene Expression Analysis

Total RNA was extracted from pooled samples of ten flies using TRIzol. cDNA was synthesized using Takara Reverse Transcriptase. Quantitative PCR (qPCR) was performed using PowerUP SYBR Green Master Mix with the primers. Results represent the average of three independent experiments. Primers used in this study were:

*Drosomycin*-F 5’-CGTGAGAACCTTTTCCAATATGAT-3’

*Drosomycin*-R 5’-TCCCAGGACCACCAGCAT-3’

*Diptericin A*-F: 5’-GCTGCGCAATCGCTTCTACT-3’

*Diptericin A*-R:5’-TGGTGGAGTGGGCTTCATG-3’

*Rpl32*-F: 5’-GACGCTTCAAGGGACAGTATCTG-3’

*Rpl32*-R: 5’-AAACGCGGTTCTGCATGAG-3’

*Pai*-F: 5’-GATCCGTTTCCTGGTCCAGTGTG-3’

*Pai*-R: 5’-CCATGAAGCTGATCGCATTGTGC-3’

*CG10332*-F: 5’-TCAGAGGCTGCACCATCAGAAGT-3’

*CG10332*-R: 5’-CCGCTTCAGAGGCTGCACC-3’

### Microbial Culturing Conditions

*Micrococcus luteus*, *Erwinia carotovora carotovora (Ecc15)*, and *Providencia burhodogranariea* strain B were cultured overnight in LB broth at 29°C. *Enterobacter cloacae*, *Escherichia coli* strain 1106, *Providencia rustigianii*, *Providencia vermicola, Providencia rettgeri,* and *Staphylococcus aureus* were cultured overnight in LB at 37°C. *Enterococcus faecalis*, *Bacillus subtilis,* and *Listeria monocytogenes* were cultured overnight in BHI at 37°C. *Metarhizium anisopliae* and *Metarhizium robertisii* were cultured on malt agar plates at 25°C for around two weeks until sporulation was observed. *Beauveria bassiana* strain R444 commercial spores were produced by Andermatt Biocontrol, products: BB-PROTEC. Bacteria were pelleted at 4000 rpm for 15 minutes at 4°C, resuspended in PBS, and diluted to the desired OD600 value indicated in the figure legends. The fungal material was filtered through glass wool using 0.05% Tween-80 to collect spores and diluted to the desired concentration (in spores/ml). All microbial strains used in this study are listed in Table S3.

### Microbiota Load Analysis

Flies were collected 30 hours post-infection and surface-sterilized with 70% ethanol. Whole-fly homogenization was performed in LB broth using a bead beater at 7200 rpm for 30 seconds. Serial dilutions of the homogenates were plated (100 µL per plate) on LB agar and incubated at 29 °C for 24 hours. Colony-forming units (CFUs) were then manually counted.

For gut microbiota load analysis, 18-day-old conventionally reared flies were collected and processed using the same procedure, except that MRS medium was used for culturing. Plates were incubated at 29 °C for 48 hours prior to CFU quantification.

### *In Vitro* Antibacterial Activity

The Paillotin peptide (45 residues) was synthesized by GenicBio with >95% purity, and Cecropin A (from silk moth) was obtained from Sigma-Aldrich (≥97% purity). Peptides were dissolved in water. Bacteria were cultured overnight and diluted to OD600 = 0.005 in LB broth, then incubated for 2 hours. Paillotin peptide and Cecropin A were diluted in LB broth to the desired concentration. Bacterial cultures (1 µL) were added to peptide solutions in a 96-well plate, and OD600 was recorded every 10 minutes using a TECAN plate reader.

### Survival Assays

Male flies (3-5 days old) were used for survival assays. The thorax of each fly was pricked with a needle (diameter ∼0.2mm) dipped in a bacterial suspension. For the fungi infection, flies were rolled in the plates of *A. fumigatus* or the commercial spores of *B. bassiana,* or were dipped into a spore solution of *M. anisopliae* or *M. robertisii*. Mortality was recorded daily. Infected flies were maintained at 29°C or 25°C and transferred to fresh vials three times per week.

### Toxin Injection

Destruxin A (Sigma) was dissolved in high-purity dimethyl sulfoxide (DMSO) and subsequently diluted with PBS to a final concentration of 8 mM. A volume of 9.2 nL of this solution was administered to flies using the Nanoject III microinjector (Drummond) [26]. Verruculogen (Abcam) was prepared in DMSO at a concentration of 1 mg/mL, and 4.6 nL of this solution was injected into the flies [31].

### Wounding experiment

Clean injury was performed with a needle sterilized (diameter ∼0.1mm) with an EtOH and PBS wash. Cuticle blackening was categorized into normal, weak, or no melanization 24h post-pricking at the injury site.

## Ethics

This work did not require ethical approval from a human subject or animal welfare committee.

## Data accessibility

All data and code used in this study are publicly available at Dryad [73].

## Declaration of AI use

We have not used AI–assisted technologies in creating this article.

## Authors’ contributions

Conceptualization: YT, XJY, MAH, BL. Data curation: YT, XJY, MAH. Funding acquisition: BL, MAH. Investigation: YT, XJY, RJJ, FS. Methodology: YT, XJY, RJJ, MAH. Project administration: BL. Validation: YT, MAH. Resources: YT, XJY, MAH. Supervision: MAH, BL. Writing – original draft: YT, BL. Writing – review & editing: MAH, BL.

## Acknowledgements.

We thank Jean-Philippe Boquette, Guiqing Liu, Samuel Rommelaere, Fanny Schupfer, Faustine Ryckebusch, and Yijie Li for experimental help and Hannah Westlake for editing. We thank Ana Marija Jakšić for providing the initial DGRP flies used to source the natural *Paillotin* mutations. We thank the Guangzhou Drosophila Resource Center (GDRC) for generating the CRISPR–Cas9 *Paillotin* knockout line. We thank Laure Menin from the EPFL ISIC Mass Spectrometry Facility for her technical assistance.

## Supplementary material

**Table S1:**
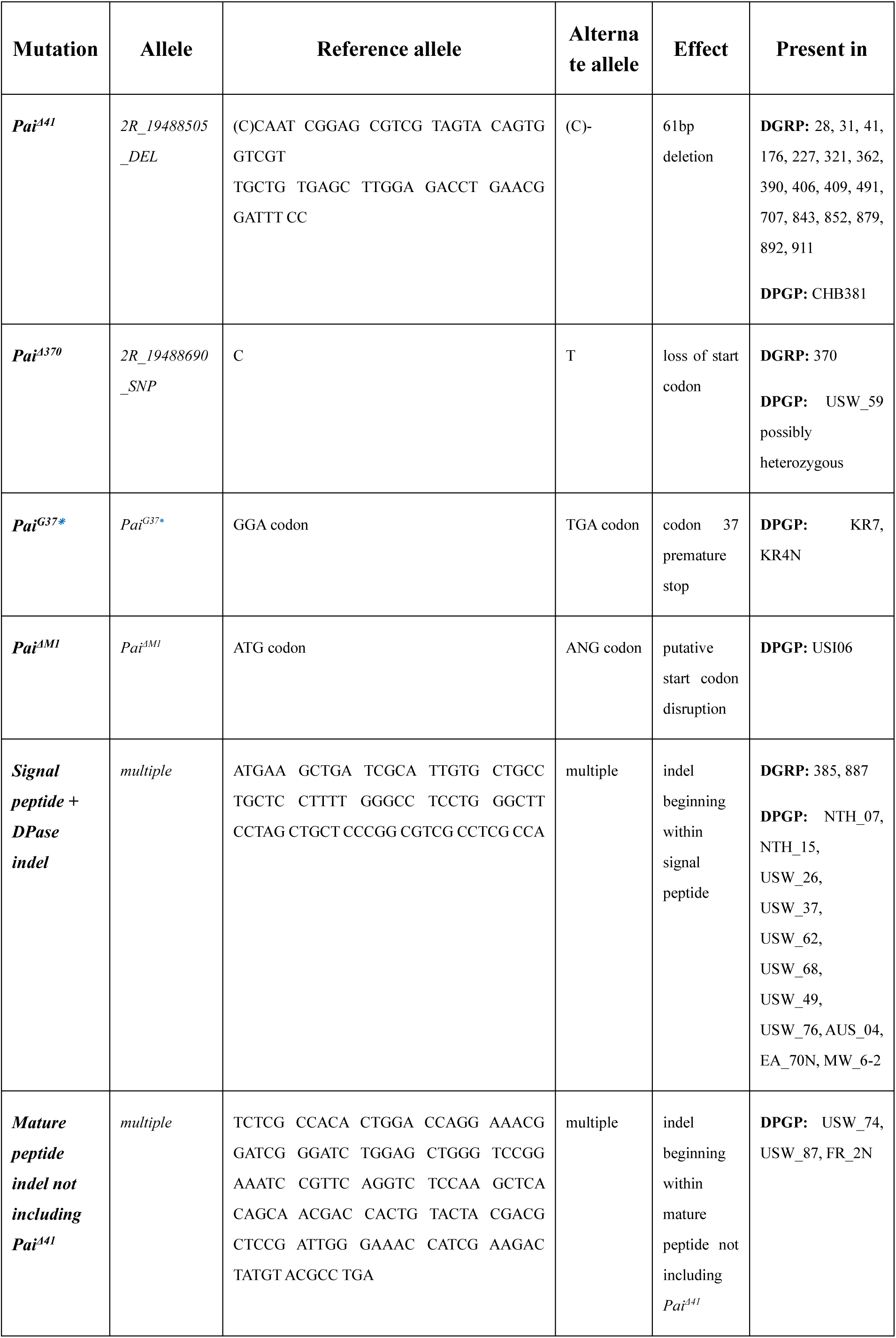
Collectively, 39 of 789 genome-sequenced fly strains, including 18 of the 205 *Drosophila* Genetic Reference Panel lines encode a loss-of-function mutation in *Paillotin*. Lines encoding *Pai* putative loss-of-function mutations are annotated, alongside the genetic description of each allele. *Pai^Δ41^* “uncertain” and “alternate” alleles per the DGRP variant call dataset are both represented by a string of ∼61 uncertain nucleotide calls (Ns) matching the pattern of allele 2R_19488505_DEL in the DPGP3 dataset (downloaded from PopFly). This suggests DGRP strains with “uncertain” allele annotations at base Dmel.R5_2R_19488505 also bear the 2R_19488505_DEL indel. Bolstered by this finding, we scanned the *Drosophila* genome nexus for strings of uncertain variant calls (Ns) that affect at least 3 contiguous codons, treating these as putative indels. This may yield an undercount by ignoring smaller indels, or an overcount by treating regions bearing multiple SNPs in linkage as indels. Nevertheless, this annotation effort draws attention to strains bearing potential loss-of-function mutations in *Paillotin* that may be relevant to future studies. DPGP indel strain summaries are grouped by the location of the first disrupted codon (within the signal or mature peptide).

**Figure S1.**
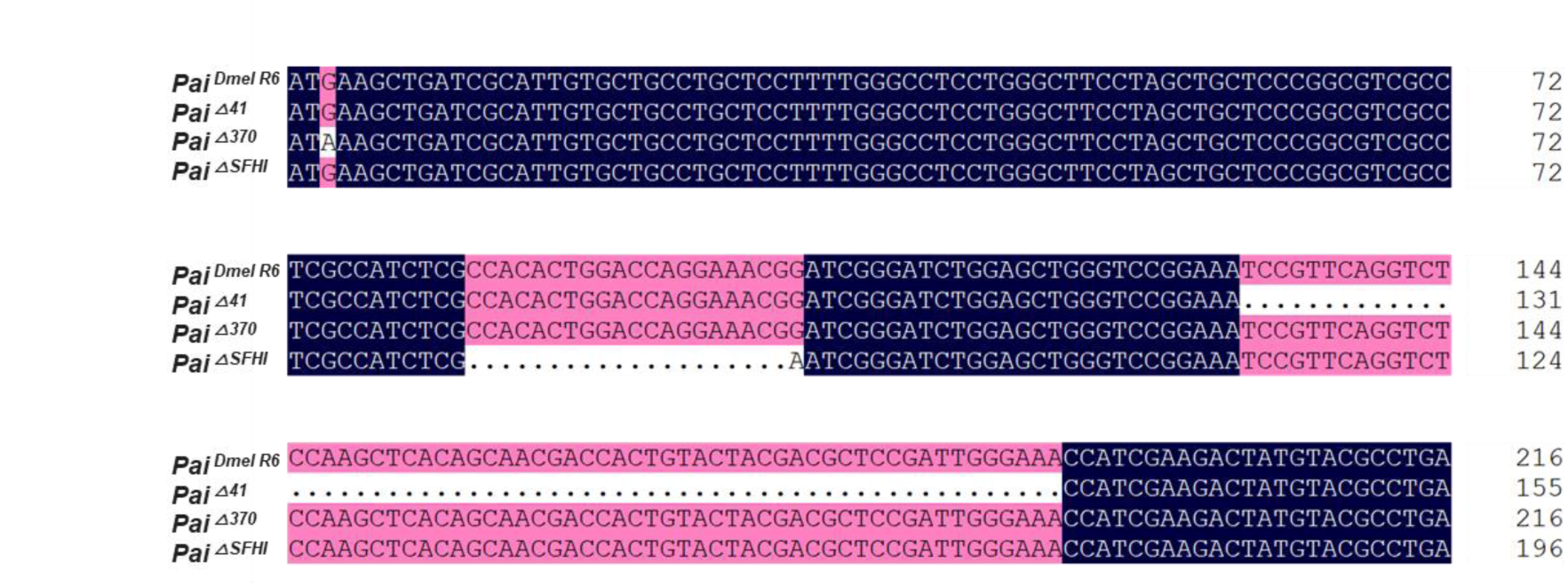
Nucleotide alignment of wild-type and mutant *Paillotin* alleles. Nucleotide sequences of wild-type *Paillotin* (*Pai^Dmel^ ^R6^*) and the three mutant alleles (*Pai*^△^*^41^*, *Pai*^△^*^370^,* and *Pai*^△^*^SFHI^*) used in this study are aligned.

**Figure S2:**
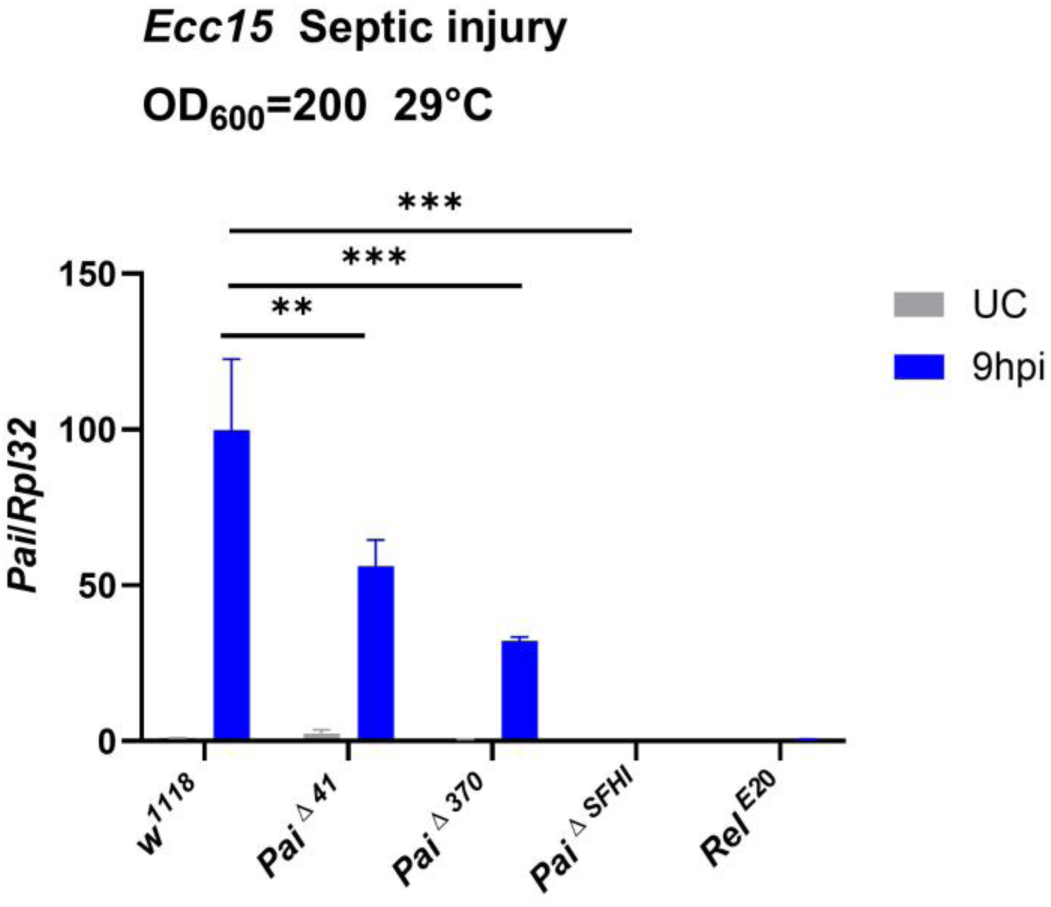
Expression of *Paillotin* in *Pai*^△^*^41^*, *Pai*^△^*^370^,* and *Pai*^△^*^SFHI^* mutations. Expression was normalized to *w^1118^* UC (unchallenged) flies set as a value of 1. The data shown in the figure are based on three independent experiments, with at least 30 flies per genotype at each time point in each experiment. Error bars represent SEM. Statistical analysis was performed using two-way ANOVA (** *p*<0.01, *** *p*<0.001).

**Figure S3:**
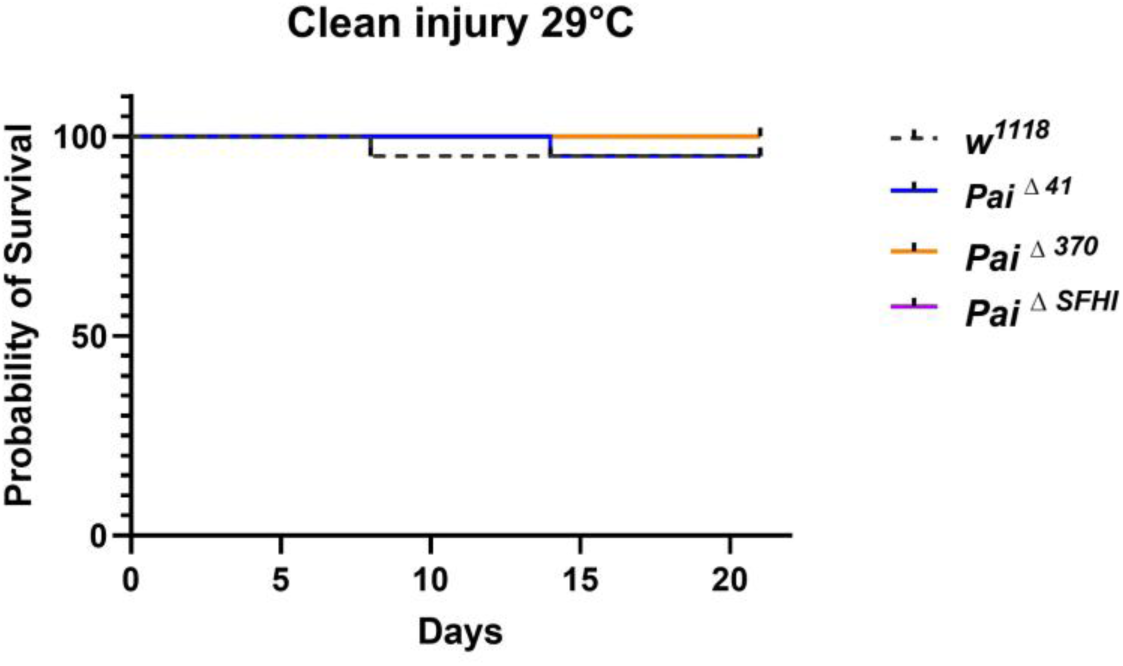
No susceptibility of *Paillotin* mutants to clean injury. The thorax of wild-type and mutants was pricked with a clean needle. A total of 80 flies were used in this experiment.

**Figure S4:**
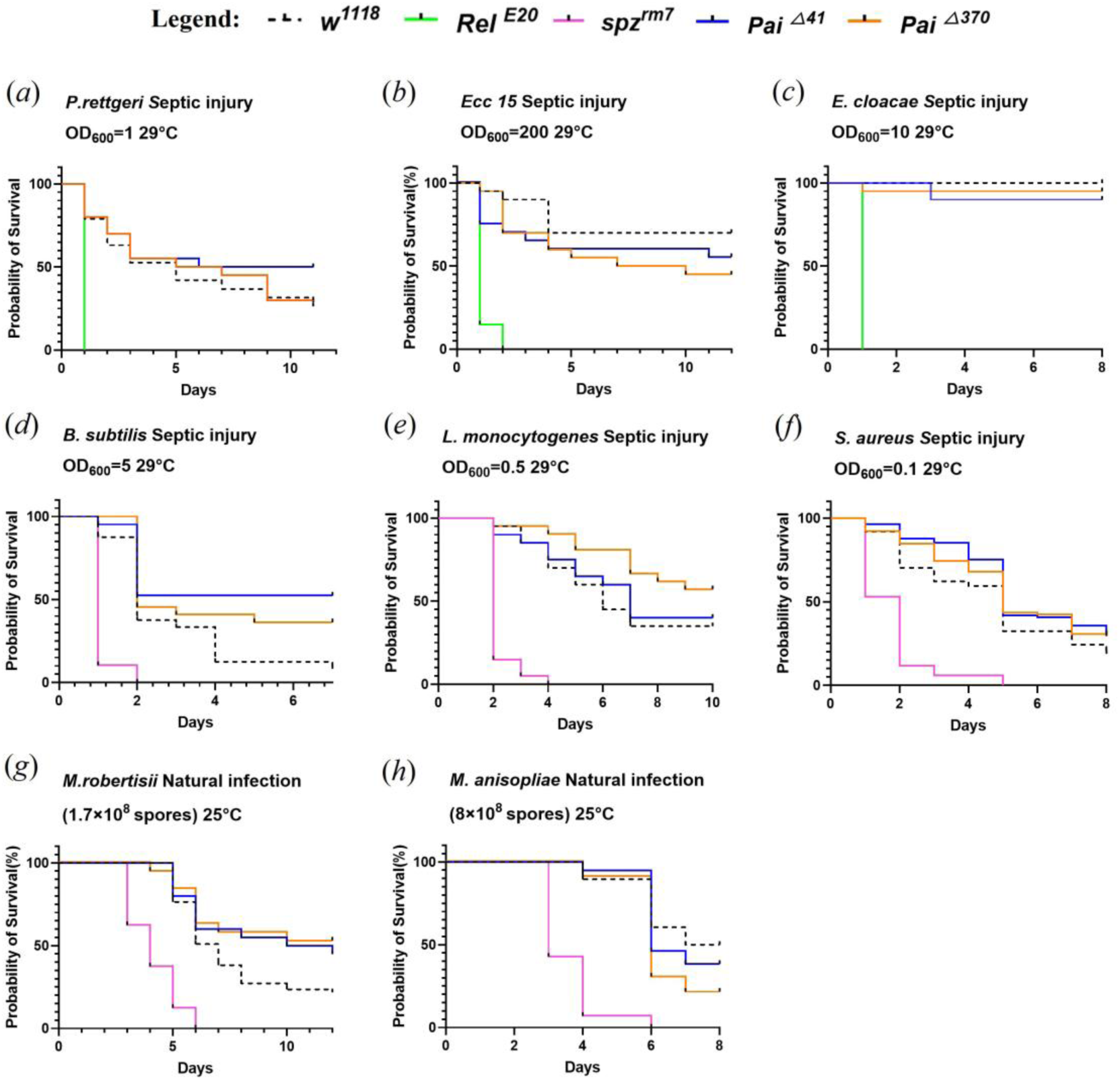
No consistent effect of *Paillotin* mutation on resistance to a broad spectrum of microbes. *w^1118^* flies were utilized as wild-type controls, while *spz^rm7^* flies lack the Toll pathway, and *Relish^E20^* flies lack the Imd pathway, were used as susceptible models for all survival experiments against infections. Tm and OD600 are indicated. Each survival assay was performed with a minimum of 80 flies.

**Figure S5:**
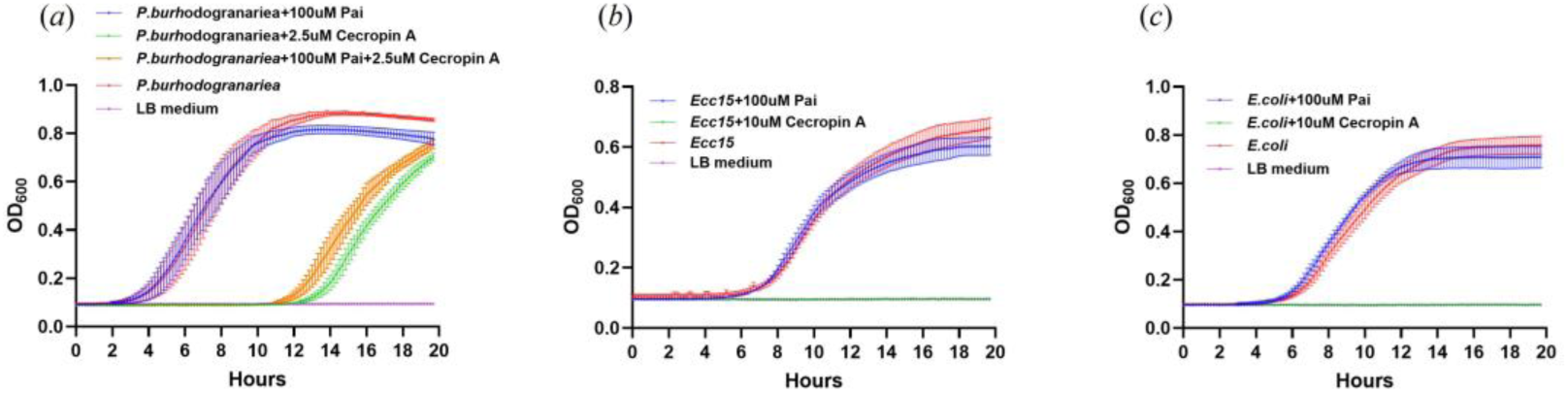
Paillotin did not show microbicidal activity *in vitro.* (*a*) The growth of *P. burhodogranariea* was monitored in the presence of the Paillotin peptide (100 µM) and/or the Cecropin A peptide (2.5 µM). The growth of *Ecc15* (*b*) and *E. coli* (*c*) was monitored in the presence of the Paillotin peptide (100 µM). Optical density (OD) values were measured every 10 minutes over 20 hours to generate bacterial growth curves. Different colored curves represent the conditions tested, as indicated in the figure legend.

**Figure S6:**
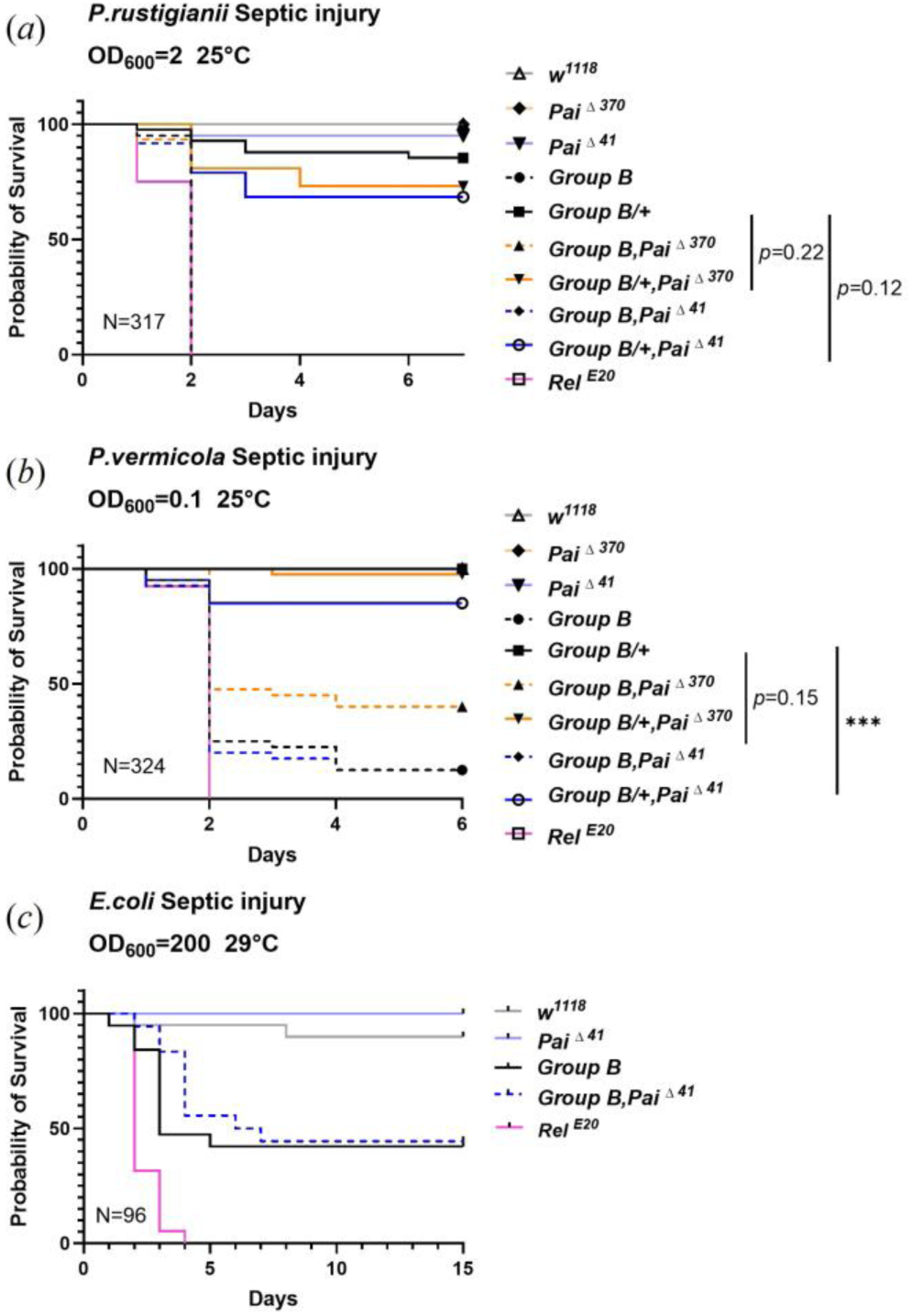
*Paillotin* shows a minor synergistic effect with *Group B* mutants against other *Providencia* species at the doses tested. (*a*) Both *Group B/+, Pai* compound mutants suffer a minor increase in mortality compared to *Group B/+* alone upon *Providencia rustigianii* infection. Although the synergistic effect between Paillotin and Group B was only marginally significant (Mutant, Cox Hazard Ratio (HR), *p-*value: *Paillotin*, HR =-0.01, *p*=0.904; *Group B*, HR =-2.8, *p*<0.001; *Paillotin*\**Group B*, HR = -0.41, *p*=0.09). (*b*) *Group B/+, Pai*^△^*^41^* mutants suffer a mortality increase compared to *Group B/+*. In addition, *Group B/+ Pai^△370^* shows no difference from *Group B/+*, but also *Group B, Pai*^△^*^370^* compound mutants survive better than *Group B* alone, indicating a complex phenotype after recombination for this infection that may mask the contribution of *Pai*^△^*^370^* at the chosen low-mortality dose (Mutant, Cox Hazard Ratio (HR), *p-*value: *Paillotin*, HR =0.15, *p*=0.14; *Group B*, HR =-2.7, *p*<0.001; *Paillotin*\**Group B*, HR = -1.13, *p*=0.03). Differences between *Pai*^△^*^41^* (truncated) and *Pai*^△^*^370^* (loss of start) may explain this difference. (*c*) No survival difference was observed for *Group B, Pai* compound mutants against *E. coli.* N=total number of flies in experiments.

**Figure S7:**
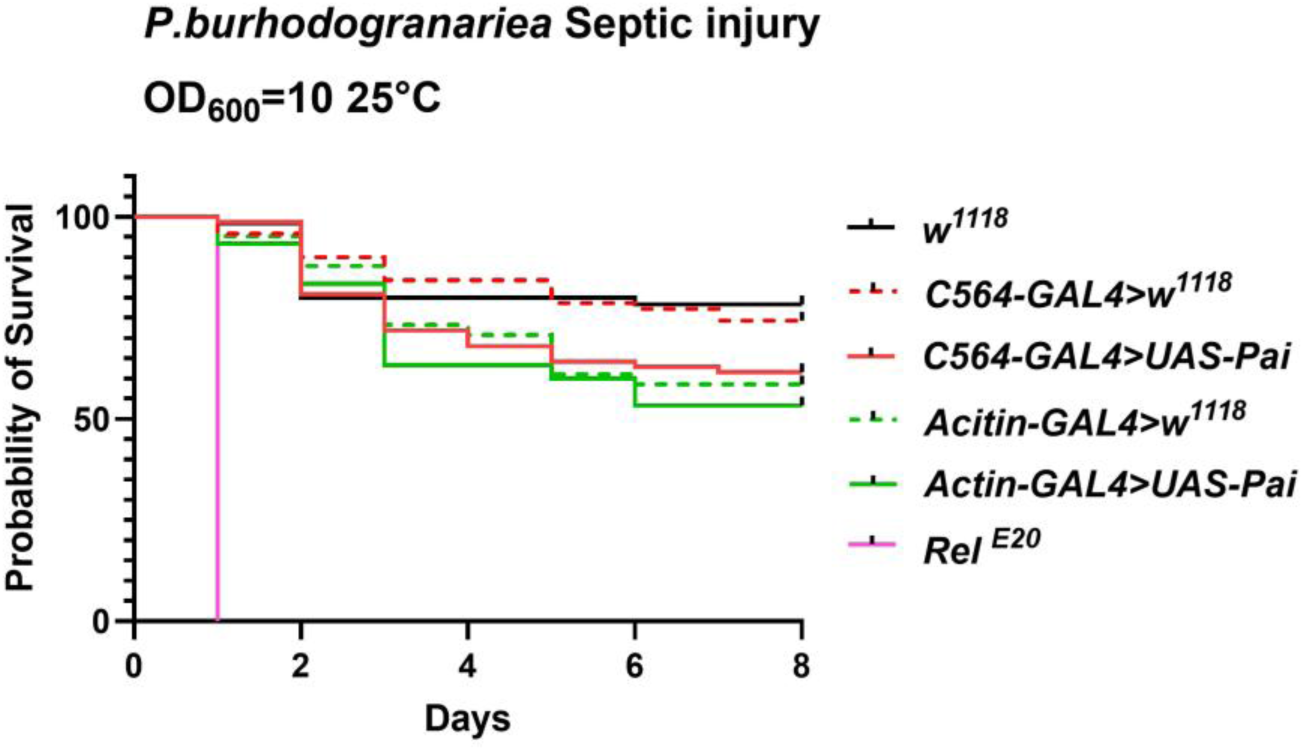
Overexpression of *Paillotin* does not confer protection against *P. burhodogranariea* infection in wild-type flies. Survival curves of flies overexpressing *Paillotin* by using *C564-Gal4* and *Actin-Gal4* drivers following septic infection with *P. burhodogranariea*. Survival assay was performed with a minimum of 120 flies.

**Figure S8:**
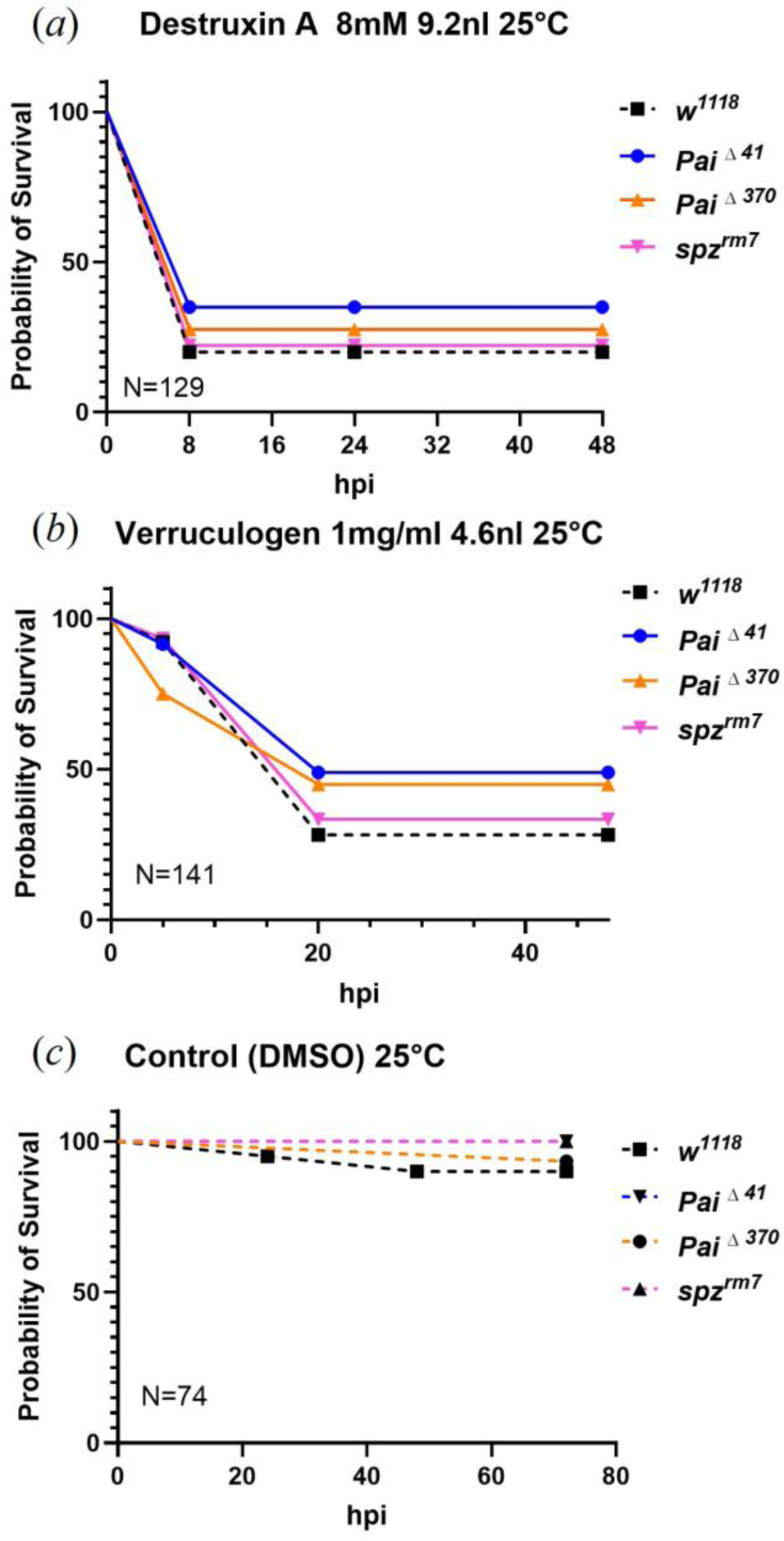
Paillotin does not contribute to protection against fungal toxins. (*a*) Survival of wild-type and *Paillotin* mutant flies following injection with 9.2 nl of 8 mM Destruxin A, a mycotoxin secreted by *M. robertsii*. (*b*) Survival of wild-type and *Paillotin* mutant flies injected with 4.6 nl of 1 mg/ml Verruculogen, a neurotoxin produced by *A. fumigatus*. (*c*) Survival of all tested fly strains following injection with DMSO as vehicle control. *spz^rm7^* flies are used as positive controls. N=total number of flies in experiments.

**Figure S9:**
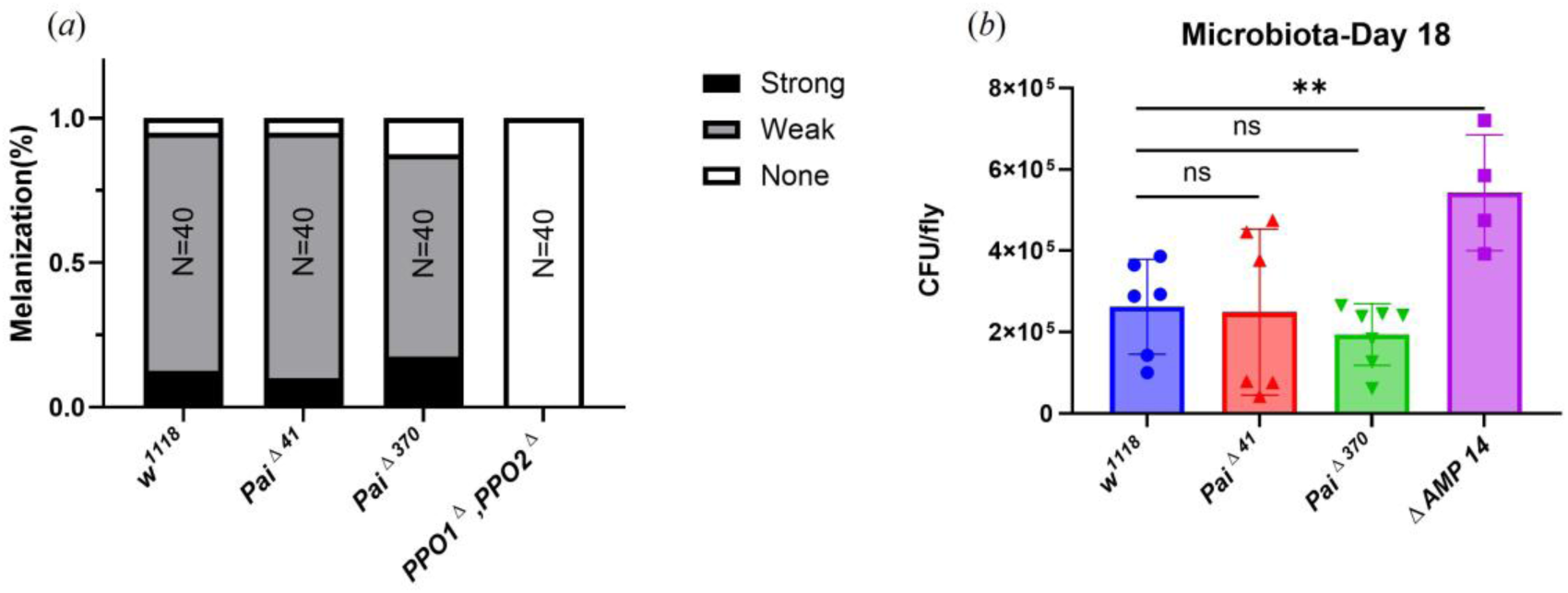
Paillotin is not involved in melanization response and microbiota regulation. (*a*) Cuticle blackening after clean injury is not impaired except in melanization-deficient flies (*PPO1^△^, PPO2^△^*). (*b*) CFUs per fly were measured for wild-type, *Paillotin* mutants, and the 14 AMP-deficient mutant (ΔAMP14). Data were analyzed using one-way ANOVA followed by Dunnett’s multiple comparisons test with *w^1118^* as the control. A significant increase in bacterial load was observed only in ΔAMP14, while no significant differences were detected between wild-type and *Paillotin* mutants. Data points represent the mean bacterial load per fly, derived from individual experiments using pools of five flies per genotype (*** *p*<0.001, n.s. = not significant, *p*>0.05).

**Table S2:** Drosophila strains used in this study.

| **Drosophila lines** | **Reference** |
| --- | --- |
| *iso; iso; Rel<sup>E20</sup>* | Hedengren et al., 1999 |
| *iso; iso; spz<sup>rm7</sup>* | Lemaitre et al., 1996 |
| *iso; Bom<sup>Δ55</sup>; iso* | Hanson et al., 2019 |
| *iso; iso; Rel<sup>E20</sup>, spz<sup>rm7</sup>* | Ryckebusch et al., 2025 |
| *iso; Pai<sup>Δ41</sup>; iso* | DGRP |
| *iso; Pai<sup>Δ370</sup>; iso* | DGRP |
| *; Pai<sup>ΔSFHI</sup>;* | Guangzhou Drosophila Resource Center |
| *; ; UAS-Pai* | This paper |
| *iso ; Group B; iso* | Hanson et al., 2019 |
| *iso ; PPO1<sup>Δ</sup>, PPO2<sup>Δ</sup>; iso* | Bingelli et al., 2014 |
| *ΔAMP 14* | Carboni et al., 2022 |
| *C564-Gal4* | BDSC |
| *Actin-Gal4* | BDSC |

**Table S3:**
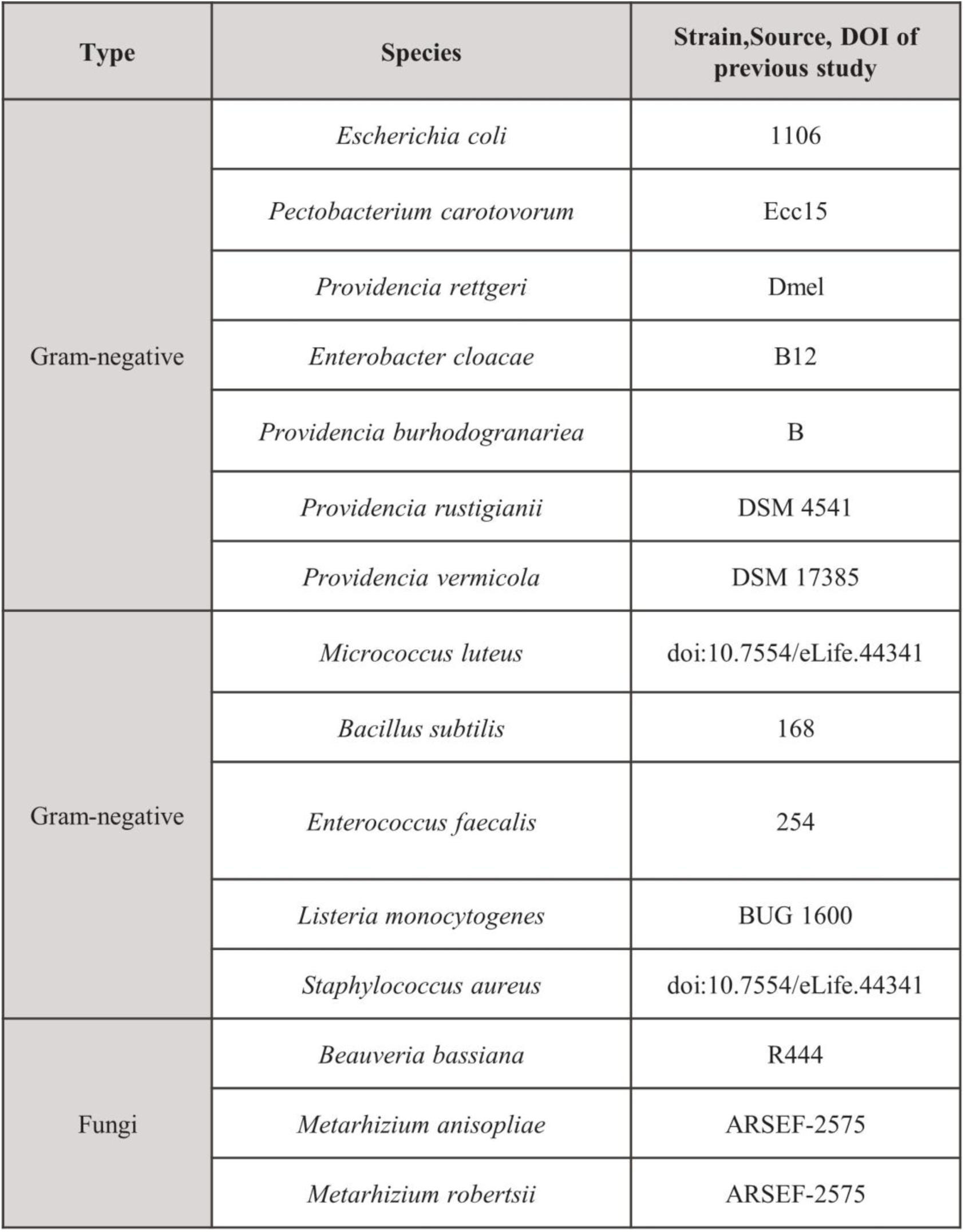
Microbial strains used in this study.

## Reference

1. Westlake H, Hanson MA, Lemaitre B. 2024 The Drosophila immunity handbook. First. EPFL Press. (doi:10.55430/6304TDIHVA01)

2. Ferrandon D, Imler J-L, Hetru C, Hoffmann JA. 2007 The Drosophila systemic immune response: sensing and signalling during bacterial and fungal infections. Nat Rev Immunol 7, 862–874. (doi:10.1038/nri2194)

3. Buchon N, Silverman N, Cherry S. 2014 Immunity in Drosophila melanogaster — from microbial recognition to whole-organism physiology. Nat Rev Immunol 14, 796–810. (doi:10.1038/nri3763)

4. Pradeu T, Thomma BPHJ, Girardin SE, Lemaitre B. 2024 The conceptual foundations of innate immunity: Taking stock 30 years later. Immunity 57, 613– 631. (doi:10.1016/j.immuni.2024.03.007)

5. De Roode JC, Lefèvre T. 2012 Behavioral Immunity in Insects. Insects 3, 789–820. (doi:10.3390/insects3030789)

6. Montanari M, Royet J. 2021 Impact of Microorganisms and Parasites on Neuronally Controlled Drosophila Behaviours. Cells 10, 2350. (doi:10.3390/cells10092350)

7. Brown KL, Hancock RE. 2006 Cationic host defense (antimicrobial) peptides. Current Opinion in Immunology 18, 24–30. (doi:10.1016/j.coi.2005.11.004)

8. Hanson MA, Lemaitre B. 2020 New insights on Drosophila antimicrobial peptide function in host defense and beyond. Current Opinion in Immunology 62, 22–30. (doi:10.1016/j.coi.2019.11.008)

9. Parvy J-P, Yu Y, Dostalova A, Kondo S, Kurjan A, Bulet P, Lemaître B, Vidal M, Cordero JB. 2019 The antimicrobial peptide defensin cooperates with tumour necrosis factor to drive tumour cell death in Drosophila. eLife 8, e45061. (doi:10.7554/eLife.45061)

10. Hirata M, Nomura T, Inoue YH. 2025 Anti-Tumor Effects of Cecropin A and Drosocin Incorporated into Macrophage-like Cells Against Hematopoietic Tumors in Drosophila mxc Mutants. Cells 14, 389. (doi:10.3390/cells14060389)

11. Marra A, Hanson MA, Kondo S, Erkosar B, Lemaitre B. 2021 *Drosophila* Antimicrobial Peptides and Lysozymes Regulate Gut Microbiota Composition and Abundance. mBio 12, e00824–21. (doi:10.1128/mBio.00824-21)

12. Kounatidis I, Chtarbanova S, Cao Y, Hayne M, Jayanth D, Ganetzky B, Ligoxygakis P. 2017 NF-κB Immunity in the Brain Determines Fly Lifespan in Healthy Aging and Age-Related Neurodegeneration. Cell Reports 19, 836–848. (doi:10.1016/j.celrep.2017.04.007)

13. E L et al. 2018 An Antimicrobial Peptide and Its Neuronal Receptor Regulate Dendrite Degeneration in Aging and Infection. Neuron 97, 125–138.e5. (doi:10.1016/j.neuron.2017.12.001)

14. Hultmark D, Steiner H, Rasmuson T, Boman HG. 1980 Insect Immunity. Purification and Properties of Three Inducible Bactericidal Proteins from Hemolymph of Immunized Pupae of *Hyalophora cecropia*. European Journal of Biochemistry 106, 7–16. (doi:10.1111/j.1432-1033.1980.tb05991.x)

15. Steiner H, Hultmark D, Engström Å, Bennich H, Boman HG. 1981 Sequence and specificity of two antibacterial proteins involved in insect immunity. Nature 292, 246–248. (doi:10.1038/292246a0)

16. Pardi A, Hare DR, Selsted ME, Morrison RD, Bassolino DA, Bach AC. 1988 Solution structures of the rabbit neutrophil defensin NP-5. Journal of Molecular Biology 201, 625–636. (doi:10.1016/0022-2836(88)90643-2)

17. Fehlbaum P, Bulet P, Michaut L, Lagueux M, Broekaert WF, Hetru C, Hoffmann JA. 1994 Insect immunity. Septic injury of Drosophila induces the synthesis of a potent antifungal peptide with sequence homology to plant antifungal peptides. Journal of Biological Chemistry 269, 33159–33163. (doi:10.1016/S0021-9258(20)30111-3)

18. Levashina EA, Ohresser S, Bulet P, Reichhart J, Hetru C, Hoffmann JA. 1995 Metchnikowin, a Novel Immune-Inducible Proline-Rich Peptide from *Drosophila* with Antibacterial and Antifungal Properties. European Journal of Biochemistry 233, 694–700. (doi:10.1111/j.1432-1033.1995.694_2.x)

19. Bulet P, Urge L, Ohresser S, Hetru C, Otvos L. 1996 Enlarged Scale Chemical Synthesis and Range of Activity of Drosocin, an O-Glycosylated Antibacterial Peptide of *Drosophila*. European Journal of Biochemistry 238, 64–69. (doi:10.1111/j.1432-1033.1996.0064q.x)

20. Ekengren S. 1999 Drosophila cecropin as an antifungal agent. Insect Biochemistry and Molecular Biology 29, 965–972. (doi:10.1016/S0965-1748(99)00071-5)

21. Rabel D, Charlet M, Ehret-Sabatier L, Cavicchioli L, Cudic M, Otvos L, Bulet P. 2004 Primary Structure and in Vitro Antibacterial Properties of the Drosophila melanogaster Attacin C Pro-domain. Journal of Biological Chemistry 279, 14853– 14859. (doi:10.1074/jbc.M313608200)

22. Clemmons AW, Lindsay SA, Wasserman SA. 2015 An Effector Peptide Family Required for Drosophila Toll-Mediated Immunity. PLoS Pathog 11, e1004876. (doi:10.1371/journal.ppat.1004876)

23. Cohen LB, Lindsay SA, Xu Y, Lin SJH, Wasserman SA. 2020 The Daisho Peptides Mediate Drosophila Defense Against a Subset of Filamentous Fungi. Front. Immunol. 11, 9. (doi:10.3389/fimmu.2020.00009)

24. Hanson MA, Cohen LB, Marra A, Iatsenko I, Wasserman SA, Lemaitre B. 2021 The Drosophila Baramicin polypeptide gene protects against fungal infection. PLoS Pathog 17, e1009846. (doi:10.1371/journal.ppat.1009846)

25. Hanson MA, Kondo S, Lemaitre B. 2022 Drosophila immunity: the Drosocin gene encodes two host defence peptides with pathogen-specific roles.Proc Biol Sci. 289, 20220773.(10.1098/rspb.2022.0773)

26. Huang J et al. 2023 A Toll pathway effector protects *Drosophila* specifically from distinct toxins secreted by a fungus or a bacterium. Proc. Natl. Acad. Sci. U.S.A. 120, e2205140120. (doi:10.1073/pnas.2205140120)

27. Hanson MA, Dostálová A, Ceroni C, Poidevin M, Kondo S, Lemaitre B. 2019 Synergy and remarkable specificity of antimicrobial peptides in vivo using a systematic knockout approach. eLife 8, e44341. (doi:10.7554/eLife.44341)

28. Carboni AL, Hanson MA, Lindsay SA, Wasserman SA, Lemaitre B. 2022 Cecropins contribute to *Drosophila* host defense against a subset of fungal and Gram-negative bacterial infection. Genetics 220, iyab188. (doi:10.1093/genetics/iyab188)

29. Hanson MA, Grollmus L, Lemaitre B. 2023 Ecology-relevant bacteria drive the evolution of host antimicrobial peptides in *Drosophila*. Science 381, eadg5725. (doi:10.1126/science.adg5725)

30. Lindsay SA, Lin SJH, Wasserman SA. 2018 Short-Form Bomanins Mediate Humoral Immunity in **Drosophila**. J Innate Immun 10, 306–314. (doi:10.1159/000489831)

31. Xu R et al. 2023 The Toll pathway mediates *Drosophila* resilience to *Aspergillus* mycotoxins through specific Bomanins. EMBO Reports 24, e56036. (doi:10.15252/embr.202256036)

32. Unckless RL, Howick VM, Lazzaro BP. 2016 Convergent Balancing Selection on an Antimicrobial Peptide in Drosophila. Current Biology 26, 257–262. (10.1016/j.cub.2015.11.063)

33. Vanha-aho L-M, Kleino A, Kaustio M, Ulvila J, Wilke B, Hultmark D, Valanne S, Rämet M. 2012 Functional Characterization of the Infection-Inducible Peptide Edin in Drosophila melanogaster. PLoS ONE 7, e37153. (doi:10.1371/journal.pone.0037153)

34. Lu M, Wei D, Shang J, Li S, Song S, Luo Y, Tang G, Wang C. 2024 Suppression of Drosophila antifungal immunity by a parasite effector via blocking GNBP3 and GNBP-like 3, the dual receptors for β-glucans. Cell Reports 43, 113642. (doi:10.1016/j.celrep.2023.113642)

35. K. Maasdorp M, Valanne S, Vesala L, Vornanen P, Haukkavaara E, Tuomela T. 2025 IbinA and IbinB regulate the Toll pathway-mediated immune response in Drosophila melanogaster. bioRxiv. (10.1101/2025.04.10.648149)

36. Goto A, Yano T, Terashima J, Iwashita S, Oshima Y, Kurata S. 2010 Cooperative Regulation of the Induction of the Novel Antibacterial Listericin by Peptidoglycan Recognition Protein LE and the JAK-STAT Pathway. Journal of Biological Chemistry 285, 15731–15738. (doi:10.1074/jbc.M109.082115)

37. Westlake H, David F, Tian Y, Krakovic K, Dolgikh A, Juravlev L, Esmangart de Bournonville T, Carboni A, Melcarne C, Shan T, Wang Y, Mu Y, Kotwal A, Pirko N, Boquete JP, Schüpfer F, Rommelaere S, Poidevin M, Liu Z, Kondo S, Ratnaparkhi GS, Chakrabarti S, Liu G, Masson F, Li X, Hanson MA, Jiang H, Di Cara F, Kurant E, Lemaitre B. 2025 Reproducibility of Scientific Claims in Drosophila Immunity: A Retrospective Analysis of 400 Publications. bioRxiv. (10.1101/2025.07.07.663442)

38. Uttenweiler-Joseph S, Moniatte M, Lagueux M, Van Dorsselaer A, Hoffmann JA, Bulet P. 1998 Differential display of peptides induced during the immune response of *Drosophila* : A matrix-assisted laser desorption ionization time-of-flight mass spectrometry study. Proc. Natl. Acad. Sci. U.S.A. 95, 11342–11347. (doi:10.1073/pnas.95.19.11342)

39. Jehle JA. 2009 André Paillot (1885–1944): His work lives on. Journal of Invertebrate Pathology 101, 162–168. (doi:10.1016/j.jip.2009.03.009)

40. Carton Y. 2019 Innate Immunity: From Louis Pasteur to Jules Hoffmann. San Diego: ISTE Press Limited - Elsevier Incorporated.

41. Robinson SW, Herzyk P, Dow JAT, Leader DP. 2013 FlyAtlas: database of gene expression in the tissues of Drosophila melanogaster. Nucleic Acids Research 41, D744–D750. (doi:10.1093/nar/gks1141)

42. Hanson MA, Lemaitre B, Unckless RL. 2019 Dynamic Evolution of Antimicrobial Peptides Underscores Trade-Offs Between Immunity and Ecological Fitness. Front. Immunol. 10, 2620. (doi:10.3389/fimmu.2019.02620)

41. Sanjay Kumar, Sanjay Kumar, Arvind Kumar Singh. 2017 Population genetics of Drosophila: Genetic Variation and Differentiation among Indian Natural Populations of Drosophila ananassae. Zoological Studies 56: e. (doi:10.6620/ZS.2017.56-01)

44. Waumans Y, Baerts L, Kehoe K, Lambeir A-M, De Meester I. 2015 The Dipeptidyl Peptidase Family, Prolyl Oligopeptidase, and Prolyl Carboxypeptidase in the Immune System and Inflammatory Disease, Including Atherosclerosis. Front. Immunol. 6. (doi:10.3389/fimmu.2015.00387)

45. Copley RR, Totrov M, Linnell J, Field S, Ragoussis J, Udalova IA. 2007 Functional conservation of Rel binding sites in drosophilid genomes. Genome Res. 17, 1327– 1335. (doi:10.1101/gr.6490707)

46. Rommelaere S, Schüpfer F, Armand F, Hamelin R, Lemaitre B. 2025 An updated proteomic analysis of *Drosophila* haemolymph after bacterial infection. Fly 19, 2485685. (doi:10.1080/19336934.2025.2485685)

47. Lack JB, Lange JD, Tang AD, Corbett-Detig RB, Pool JE. 2016 A Thousand Fly Genomes: An Expanded*Drosophila*Genome Nexus. Mol Biol Evol 33, 3308–3313. (doi:10.1093/molbev/msw195)

48. Mackay TFC et al. 2012 The Drosophila melanogaster Genetic Reference Panel. Nature 482, 173–178. (doi:10.1038/nature10811)

49. Scott MG, Davidson DJ, Gold MR, Bowdish D, Hancock REW. 2002 The Human Antimicrobial Peptide LL-37 Is a Multifunctional Modulator of Innate Immune Responses. The Journal of Immunology 169, 3883–3891. (doi:10.4049/jimmunol.169.7.3883)

50. Dong X et al. 2022 Keratinocyte-derived defensins activate neutrophil-specific receptors Mrgpra2a/b to prevent skin dysbiosis and bacterial infection. Immunity 55, 1645–1662.e7. (doi:10.1016/j.immuni.2022.06.021)

51. Samakovlis C, Kimbrell DA, Kylsten P, Engström A, Hultmark D. 1990 The immune response in Drosophila: pattern of cecropin expression and biological activity. The EMBO Journal 9, 2969–2976. (doi:10.1002/j.1460-2075.1990.tb07489.x)

52. De Gregorio E. 2002 The Toll and Imd pathways are the major regulators of the immune response in Drosophila. The EMBO Journal 21, 2568–2579. (doi:10.1093/emboj/21.11.2568)

53. Tzou P, Reichhart J-M, Lemaitre B. 2002 Constitutive expression of a single antimicrobial peptide can restore wild-type resistance to infection in immunodeficient *Drosophila* mutants. Proc. Natl. Acad. Sci. U.S.A. 99, 2152–2157. (doi:10.1073/pnas.042411999)

54. Hanson MA. 2024 When the microbiome shapes the host: immune evolution implications for infectious disease. Phil. Trans. R. Soc. B 379, 20230061. (doi:10.1098/rstb.2023.0061)

55. Maiti S, Patro S, Purohit S, Jain S, Senapati S, Dey N. 2014 Effective Control of Salmonella Infections by Employing Combinations of Recombinant Antimicrobial Human β-Defensins hBD-1 and hBD-2. Antimicrob Agents Chemother 58, 6896– 6903. (doi:10.1128/aac.03628-14)

56. Myhrman E, Håkansson J, Lindgren K, Björn C, Sjöstrand V, Mahlapuu M. 2013 The novel antimicrobial peptide PXL150 in the local treatment of skin and soft tissue infections. Appl Microbiol Biotechnol 97, 3085–3096. (doi:10.1007/s00253-012-4439-8)

57. Rivas-Santiago B, Rivas Santiago CE, Castañeda-Delgado JE, León–Contreras JC, Hancock REW, Hernandez-Pando R. 2013 Activity of LL-37, CRAMP and antimicrobial peptide-derived compounds E2, E6 and CP26 against Mycobacterium tuberculosis. International Journal of Antimicrobial Agents 41, 143–148. (doi:10.1016/j.ijantimicag.2012.09.015)

58. Björn C, Håkansson J, Myhrman E, Sjöstrand V, Haug T, Lindgren K, Blencke H- M, Stensvåg K, Mahlapuu M. 2012 Anti-infectious and anti-inflammatory effects of peptide fragments sequentially derived from the antimicrobial peptide centrocin 1 isolated from the green sea urchin, Strongylocentrotus droebachiensis. AMB Expr 2. (doi:10.1186/2191-0855-2-67)

59. Zdybicka-Barabas A, Mak P, Klys A, Skrzypiec K, Mendyk E, Fiołka MJ, Cytryńska M. 2012 Synergistic action of Galleria mellonella anionic peptide 2 and lysozyme against Gram-negative bacteria. Biochimica et Biophysica Acta (BBA) - Biomembranes 1818, 2623–2635. (doi:10.1016/j.bbamem.2012.06.008)

60. Rahnamaeian M, Cytryńska M, Zdybicka-Barabas A, Vilcinskas A. 2016 The functional interaction between abaecin and pore-forming peptides indicates a general mechanism of antibacterial potentiation. Peptides 78, 17–23. (doi:10.1016/j.peptides.2016.01.016)

61. Rahnamaeian M et al. 2015 Insect antimicrobial peptides show potentiating functional interactions against Gram-negative bacteria. Proc. R. Soc. B. 282, 20150293. (doi:10.1098/rspb.2015.0293)

62. Zanchi C, Johnston PR, Rolff J. 2017 Evolution of defence cocktails: Antimicrobial peptide combinations reduce mortality and persistent infection. Molecular Ecology 26, 5334–5343. (doi:10.1111/mec.14267)

63. Engström P, Carlsson A, Engström A, Tao ZJ, Bennich H. 1984 The antibacterial effect of attacins from the silk moth Hyalophora cecropia is directed against the outer membrane of Escherichia coli. EMBO J 3, 3347–3351. (doi:10.1002/j.1460-2075.1984.tb02302.x)

64. Tyler-Cross R, Schirch V. 1991 Effects of amino acid sequence, buffers, and ionic strength on the rate and mechanism of deamidation of asparagine residues in small peptides. Journal of Biological Chemistry 266, 22549–22556. (doi:10.1016/s0021-9258(18)54607-x)

65. Alkotaini B, Anuar N, Kadhum AAH, Sani AAA. 2013 Detection of secreted antimicrobial peptides isolated from cell-free culture supernatant of *Paenibacillus alvei* AN5. Journal of Industrial Microbiology and Biotechnology 40, 571–579. (doi:10.1007/s10295-013-1259-5)

66. Samakovlis C, Kylsten P, Kimbrell DA, Engström A, Hultmark D. 1991 The andropin gene and its product, a male-specific antibacterial peptide in Drosophila melanogaster. The EMBO Journal 10, 163–169. (doi:10.1002/j.1460-2075.1991.tb07932.x)

63. Ryckebusch F, Tian Y, Rapin M, Schupfer F, Hanson M, Lemeitre B. 2025 Layers of immunity: Deconstructing the Drosophila effector response. *bioRxiv*. (doi: 10.1101/2025.03.28.645959)

68. Chen J-S, Tsaur S-C, Ting C-T, Fang S. 2022 Dietary Utilization Drives the Differentiation of Gut Bacterial Communities between Specialist and Generalist Drosophilid Flies. Microbiol Spectr 10, e01418–22. (doi:10.1128/spectrum.01418-22)

69. Kim BY et al. 2021 Highly contiguous assemblies of 101 drosophilid genomes. eLife 10, e66405. (doi:10.7554/eLife.66405)

70. Kearse M et al. 2012 Geneious Basic: An integrated and extendable desktop software platform for the organization and analysis of sequence data. Bioinformatics 28, 1647–1649. (doi:10.1093/bioinformatics/bts199)

71. Ferreira ÁG, Naylor H, Esteves SS, Pais IS, Martins NE, Teixeira L. 2014 The Toll-Dorsal Pathway Is Required for Resistance to Viral Oral Infection in Drosophila. PLoS Pathog 10, e1004507. (doi:10.1371/journal.ppat.1004507)

72. Levy F, Rabel D, Charlet M, Bulet P, Hoffmann JA, Ehret-Sabatier L. 2004 Peptidomic and proteomic analyses of the systemic immune response of Drosophila. Biochimie 86, 607–616. (doi:10.1016/j.biochi.2004.07.007)

73. Tian Y, Yue XJ, Jiao RJ, Hanson M, Lemaitre B. 2025 Data from: Functional characterization of Paillotin: an immune peptide regulated by the Imd pathway with pathogen-specific roles in Drosophila immunity. Dryad Digital Repository. (10.5061/dryad.f4qrfj77k)

